# VARS1 fuels prostate cancer evolution via codon-selective translational rewiring

**DOI:** 10.64898/2025.12.09.693333

**Authors:** Qinju He, Rongrong He, Yingying Li, Miaomiao Xu, Shuchao Ren, Cheng Zou, Zhenyu Li, Wenchao Li, Yuanzhen Zhang, Lanxin Hu, Bin Xu, Baobing Zhao, Zhipeng Zhou, Dingxiao Zhang

**Author notes:** These authors contributed equally (Qinju He, Rongrong He, Yingying Li, Miaomiao Xu, Shuchao Ren, Cheng Zou). **Note:** Q. He, R. He, Y. Li, M. Xu, S. Ren and C. Zou contributed equally to this article. **Corresponding Authors**: Dingxiao Zhang, Hunan University, Changsha 410082, China.; Zhipeng Zhou, Huazhong Agricultural University, Wuhan 430070, China.; Baobing Zhao, Shandong University, Jinan 250012, China.; Bin Xu, Southeast University; Nanjing 210009, China.

## Abstract

Cancers exhibit translatomic alterations, but little is known about the drivers that control translational dysregulation and can also be exploited therapeutically in prostate cancer (PCa). By systematic interrogating a group of genes associated with transfer RNA (tRNA) biology (termed as tRNA biogenesis), here we establish tRNA biogenesis as an overall oncogenic pathway in, and identify valyl-tRNA synthetase (VARS1) as the key underlying driver of, PCa progression. Targeting VARS1 reduces the charged levels of valine tRNAs, inhibits global translation and suppresses aggressive PCa both in vitro and in vivo. Surprisingly, knocking down VARS1 does not preferentially impact on translation of valine-rich transcripts, but instead switches the usage of PCa-preferred GTA and GTT codons to GTC and GTC codons, which are optimal in normal prostates. Mechanistically, overexpressed VARS1 in PCa selectively accelerates the translation of genes with high GTA and GTT codon content that are functionally tied to cell mitosis and cancer-promoting pathways. Dietary valine restriction (VR) reduces global translation and slows the growth of both AR^+^ and AR^—^xenograft models. We have also developed a VARS1 inhibitor that suppresses autochthonous prostate tumours by targeting its aminoacylation activity. Altogether, our studies indicate that VARS1 acts as an oncogene promoting PCa progression through codon-selective translational rewiring, and therefore represents a therapeutic target susceptible to dietary VR and small molecule therapy.

## INTRODUCTION

Androgen deprivation therapy (ADT) is the main management strategy for prostate cancer (PCa), the most prevalent form of male cancer worldwide [1]. Despite initial effectiveness, most patients treated with ADT invariably develop castration-resistant PCa (CRPC), currently a lethal form of the disease [2]. Second-line regimens aiming to further block androgen receptor (AR) function (using enzalutamide) and/or block adrenal androgen biosynthesis (using abiraterone) have been developed for CRPC, but, unfortunately, patients still experience disease recurrence with a shorter interval due to acquired resistance [3, 4]. Consequently, PCa still causes a significant mortality among men globally, mainly owing to our incomplete understanding of the mechanisms governing tumour progression, especially treatment resistance and subsequent maintenance of CRPC [5, 6]. Therefore, identifying novel therapeutic targets and developing new clinical treatment strategies are imperative to combat aggressive PCa.

Centered on central-dogma of molecular biology (from DNA to messenger RNA (mRNA) to protein), studies in the past decades, owing to the global efforts of applying high-throughput next-generation sequencing technologies in clinic (typified by The Cancer Genome Atlas (TCGA) program [7, 8]), have now well-established alterations in cancer genome and/or transcriptome as pivotal drivers in prostate tumourigenesis. Although less efforts, relatively, have been made to investigate the proteome compared with extensively studied genome and epigenome of PCa, a few recent large-scale proteomic profiling studies on clinical primary PCa (pri-PCa) samples from both Western [9] and Chinese [10–12] patients have firmly confirmed proteomic dysregulation as a hallmark of PCa development. Importantly, the mRNA abundance changes explain globally only about 10% of protein abundance variability [9], highlighting that abnormality in mRNA translation is the determinant of shaping aberrant PCa proteome. Broadly, mounting evidence indicates that mRNA translational reprogramming drives protumourigenic programs [13]. For instance, translation initiation, the first and often the rate-limiting step in mRNA translation, is frequently hijacked by oncogenic signaling pathways [13, 14]. Previously, a report in animal models showed that prostatic AR can directly upregulate the translation inhibitor 4EBP1 transcriptionally, resulting a decreased activity of translation initiation and so as the protein synthesis [15]. This study therefore proposed that AR-low PCa might be more vulnerable to disruption of the eIF4F initiation complex. Nevertheless, in the essence of translational control, mechanisms that dictate the global proteomic dysregulation during PCa evolution remain largely unknown.

Integral to translation, aminoacylated transfer RNA (tRNA), which is catalyzed by aminoacyl-tRNA synthetase (aaRS) via covalent ligation of each amino acid to cognate isoacceptor tRNAs, is a key substrate in protein synthesis [13]. Altered tRNA dynamics has been reported to play a vital role in the translation of cancer genomes [16]. In this study, we refer to the genes involved in different aspects of tRNA biogenesis (e.g., aaRS, tRNA modification enzymes and others), broadly, as tRNA-biology regulatory genes (TBGs) (Table S1). As expected, recent studies also revealed that specific aaRS can either promote or suppress tumourigenesis [17]. The leucyl-tRNA synthetase (LARS) is repressed in expression in breast cancer transformation and functions as a tumour suppressor by regulating codon-dependent translation of growth suppressive genes [18]. The bifunctional glutamyl-prolyl-tRNA synthetase (EPRS1) exerts oncogenic role in multiple myeloma (MM) and is associated with poor patient prognosis [19]. As such, blockade of aaRS is nominated as potential anticancer therapeutics. For example, NCP26, an ATP-competitive EPRS1 inhibitor, demonstrates significant anti-tumour activity in MM [19]. Another study also reported that inhibitors of EPRS1 prevent fibrosis by blocking the biosynthesis of collagen, a proline-rich protein [20].

Here we systematically analyze the dysregulated TBGs and their clinical relevance, and identify overexpressed valine aminoacyl-tRNA synthetase (VARS1) as a key node shaping PCa-specific proteome. By combining ribosome profiling, polysome profiling followed by RNA sequencing (RNA-seq) of polysome-bound mRNAs (polysome-seq), ribosome footprint sequencing (Ribo-seq) and quantitative proteomics, we demonstrate that VARS1 drives PCa evolution through the translation of valine-enriched transcripts in a codon-switch manner. Therapeutically, we propose dietary valine restriction (VR) and small-molecule VARS1 inhibitors as actionable strategies to target aggressive PCa.

## RESULTS

### 1. Global dysregulation in tRNA biogenesis associates with PCa progression

To determine whether tRNA biogenesis is deregulated in human PCa, we ***first*** surveyed the expression of a curated set of 105 TBGs (Fig. 1A and Table S1) during PCa development and progression. Comparative analyses of either bulk (Fig. S1A) or paired (Fig. 1B) tumours (T) versus (vs.) normal (N) tissues in curated [7] and uncurated (pan-cancer) TCGA cohorts indicated that a group of TBGs were misexpressed in pri-PCa, with almost all genes being upregulated in tumours (Fig. 1C and Table S2). Strikingly, abnormality in TBGs expression became markedly exacerbated when tumours reached CRPC stage (Fig. 1D and Fig. S1B). Notably, more TBGs were also upregulated in CRPC relative to pri-PCa (Fig. 1E). Aggregately, we observed 55 (52.4%) genes deregulated in any comparison by a cutoff of foldchange (FC) difference ≥ |1.5| and false discovery rate (FDR) < 0.05, and 86 (81.9%) genes by FC ≥ |1.3| and FDR<0.05 (Table S2). ***Second***, to further explore the clinical relevance of tRNA biogenesis, we assessed the prognostic values of dysregulated TBGs in patient’s outcome. Using the curated TCGA [7] and SU2C CRPC [8] cohorts as representatives, we found that 80% and 48% of dysregulated TBGs in pri-PCa and CRPC, respectively, were unfavorable genes whose higher expression predicted poor patient survival (Fig. 1F). No TBGs in pri-PCa and only one TBG in CRPC were favorable genes whose higher expression correlated with better patient survival (Fig. S1C). ***Third***, we next systematically surveyed the mutational landscape of 105 TBGs in 9 large-scale clinical datasets in cBioportal [21], which were categorized as pri-PCa and CRPC datasets. Globally, CRPC (vs. pri-PCa) displayed a higher frequency in genomic alterations, especially amplification, in TBGs (Fig. 1G). Examination of the mutational landscape of top 16 altered TBGs across all datasets (Fig. S1D) or in the representative pri-PCa (Fig. S1E) and CRPC (Fig. S1F) datasets showed that most individual TBGs were mutated at low frequency (<5%). However, in combination, TBGs collectively represent a frequently mutated pathway in PCa, as 52% of patients with PCa (regardless of cancer stages) harbor at least one mutation of one TBG (Fig. 1G). Importantly, classification of patients with or without mutations in TBGs showed that the mutant group displayed a markedly worse clinical outcome (Fig. 1H and Fig. S1G), suggesting a potential protumourigenic role for the deregulated tRNA pathway. ***Fourth***, to further establish a functional dependency of TBGs in PCa, we mined a large-scale CRISPR-Cas9 gene essentiality screen deposited in Cancer Dependency Map (https://depmap.org/portal/), and then overlapped the TBGs with top 1500 essential genes found in each PCa lines. On average, 33 out of 105 genes in CRISPR screens were found essential, comparably, in both androgen-sensitive AR^+^ and CRPC-like AR^−^ cell lines (Fig. 1I), highlighting their biological importance. Interestingly, overlap of essential TBGs found in any of the PCa lines (n=54) with upregulated TBGs (n=39 by FC1.5 and n=64 by FC1.3) identified in any of the paired comparisons between different disease entities (Fig. 1C and 1E) indicated strongly that essential TBGs are mainly upregulated ones (Fig. S1H). Altogether, these results suggest that altered tRNA biogenesis augments along PCa evolution and potentially functions as an oncogenic pathway.

**Fig. 1.**
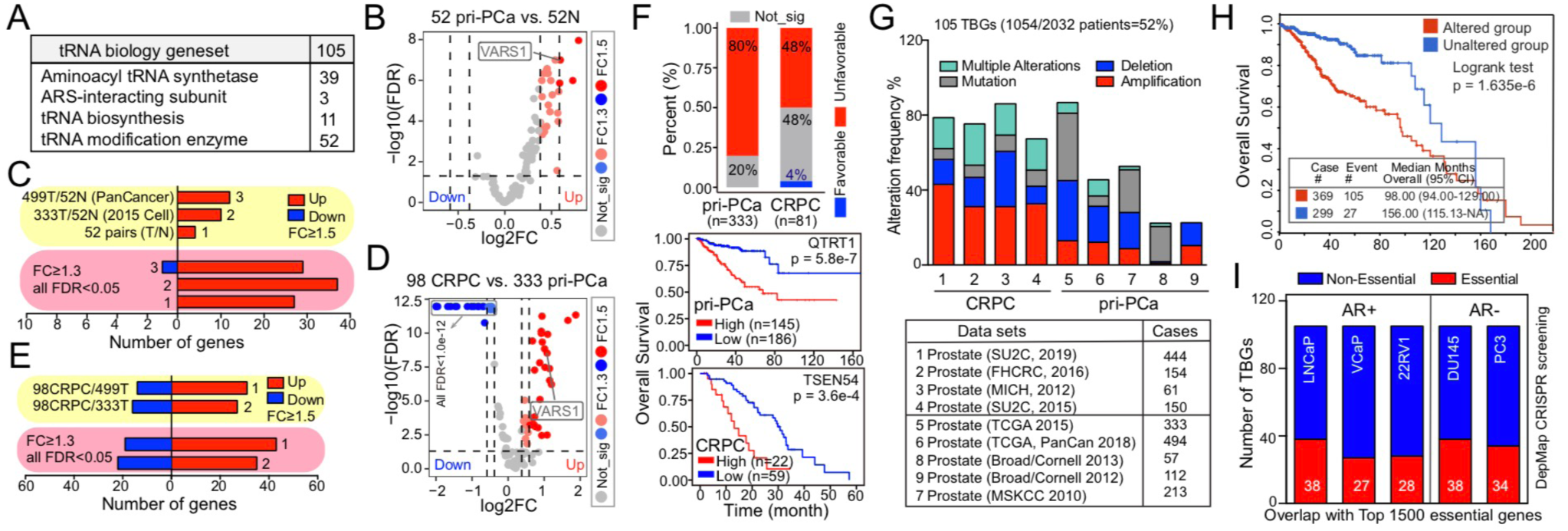
Clinical evidence that cumulative dysregulation in tRNA biogenesis associates with PCa progression. **A.** Classification of tRNA biology regulatory genes (TBGs, also see Table S1 for detail). **B.** Volcano plot showing the differentially expressed TBGs identified in 52 pairs of pri-PCa vs. normal/benign tissues with indicated statistical cutoffs. **C.** Numbers of genes showing significant up- or down-regulation in indicated comparisons in the TCGA cohorts. **D.** Volcano plot showing the differentially expressed TBGs identified in CRPC vs. pri-PCa comparison with indicated statistical cutoffs. **E.** Numbers of genes showing significant up- or down-regulation in indicated comparisons in the SU2C-CRPC cohort. **F.** Percentage of differentially expressed TBGs showing favorable (blue) and unfavorable (red) prognostic effects in TCGA and CRPC cohorts (top). Kaplan–Meier plots for representative unfavorable genes associated with patient overall survival in indicated cohorts (bottom). **G.** Bar plots illustrating the cumulative aberration frequencies of all 105 TBGs combined across all cohorts. **H.** Kaplan–Meier plot showing groups with (vs. without) genomic mutations in 105 TBGs being significantly associated with a worse patients’ overall survival (OS). **I.** Numbers of TBGs displaying functional essentiality in five PCa cell lines as revealed by DepMap CRISPR screening project (https://depmap.org/portal/). Top 1500 essential genes discovered in each cell line were used for overlapping.

### 2. Upregulation of VARS1 in PCa correlates with worse survival

To identify key TBGs that might drive PCa progression, we overlapped upregulated TBGs in 5 different pair-comparisons as indicated (Fig. 2A), and identified VARS1 as the only and central gene overlapping. Examination of *VARS1* mRNA levels in curated [7] (Fig. 2B) and uncurated (Fig. S2A) pri-PCa TCGA cohorts and in many other datasets recorded in the PCaDB database (including Taylor and other 10 datasets) [22] indicated that, *VARS1* was uniformly overexpressed in pri-PCa compared to N or prostatic intraepithelial neoplasia (PIN) tissues (Fig. 2B and Fig. S2B). Strikingly, *VARS1* mRNA was further increased in clinical CRPC vs. pri-PCa samples (Fig. 2B and Fig. S2A). Consistently, immunohistochemistry (IHC) analysis in clinical specimens encompassing the whole spectrum of PCa evolution revealed that VARS1 protein levels were higher in pri-PCa vs. N tissues and further elevated in CRPC and metastases relative to pri-PCa (Fig. 2C). Notably, the *VARS1* expression was positively associated with aggressive clinical features such as increased pathological Gleason score (GS) and tumour stages, and with decreased overall survival (OS) in both curated (Fig. 2D) and uncurated (Fig. S2C) TCGA cohorts. Moreover, Kaplan–Meier analysis in other 4 pri-PCa datasets also unraveled a negative prognostic value of *VARS1* expression for predicting patients’ OS (Fig. S2D). Importantly, an adverse correlation of *VARS1* mRNA with CRPC patients’ OS [23] was also observed (Fig. 2E).

**Fig. 2.**
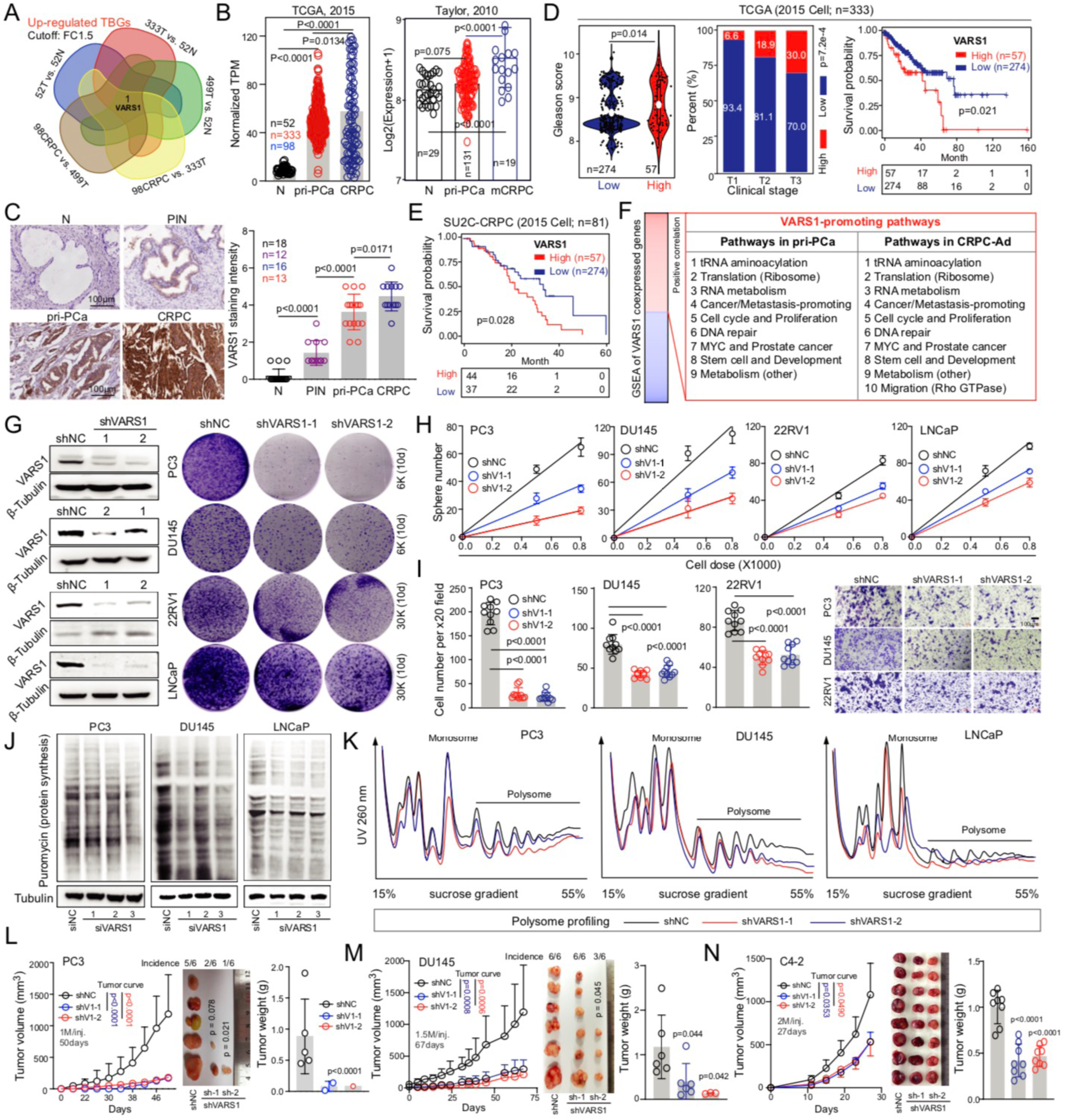
VARS1 is essential for PCa maintenance and translation. **A.** Venn diagram showing VARS1 as the only gene overlapped in upregulated TBGs identified in indicated five different comparisons. **B.** Continuous overexpression of VARS1 at mRNA level during PCa development (pri-PCa vs. N) and progression (CRPC vs. pri-PCa). **C.** IHC analysis of VARS1 protein in a PCa TMA. Representative IHC images of VARS1 in normal, PIN, pri-PCa and CRPC tissues (left) with quantification data (right) are shown. **D.** Comparison of GS (left), tumour stage (middle) and patients’ OS (right) showing the VARS1^high^ group being more clinically aggressive in the curated TCGA cohort. **E.** High VARS1 mRNA levels correlate with a worse patient OS in the indicated CRPC cohort. **F.** GSEA of genes co-expressed with VARS1 in both curated TCGA pri-PCa and CRPC cohorts. **G.** Western blot (left) and colony formation analysis (right) showing the knockdown (KD) efficiency of shRNAs targeting VARS1 in a panel of PCa cells, and the impact of VARS1 depletion on cancer cell clonal development (cell dose indicated), respectively. **H** and **I**. Knocking down VARS1 reduces sphere formation (**H**) and migration (**I**) in indicated PCa cell lines. Note that the aggressiveness generally increased from AR^+^ LNCaP to AR^—^ DU145 to stem-like PC3 and VARS1 KD affected PC3 more prominently than AR^+^ cells in these assays. Quantifications (left) and representative low magnification images (right) and are shown for cell migration assay (**I**). Data represent the means ± SD from cell number counting of at least 10 random high magnification (20X) images. **J.** Puromycin incorporation assay showing a visible reduction of total protein synthesis rate in a panel of PCa cells upon VARS1 KD. **K.** Polysome traces from gradient fractionation and polysome profiling showing reduced global translation in VARS1-depleted PCa cells compared to control. Peaks corresponding to monosome and polysome are labeled. **L-N**. Targeting VARS1 inhibits tumour regeneration and growth of AR^—^ PC3 (n = 6 for each group; **L**), DU145 (n = 6 for each group; **M**) and AR^+^ C4-2 (n = 8 for each group; **N**) cells. Shown are the tumour growth curves (left), tumour incidence and endpoint tumour images (middle), and tumour weight (right) for indicated models. Data represent mean ± SD, and the *P*-values for tumour incidence, tumour growth curve and weight were determined using χ^2^ test and two-tailed unpaired Student’s t-test, respectively.

To further gain mechanistic insights underpinning VARS1’s function, we performed a gene-coexpression analysis, coupled with gene set enrichment analysis (GSEA), to identify biological gene signatures and pathways that may be regulated by VARS1. Both TCGA pri-PCa [7] and CRPC [23] cohorts were examined and similar repertoires of biological pathways were identified (Fig. 2F and Table S3), indicating conserved roles of VARS1 in both treatment-naïve and treatment-resistant PCa. Globally, genes co-expressed with *VARS1* were prominently enriched in pathways of tRNA aminoacylation, translation and RNA metabolism (such as splicing and transcription) (Fig. 2F and Fig. S2E), in line with VARS1’s intrinsic roles in these pathways and also validating our analytic pipeline. Notably, pathways tied to cancer-promoting functions (e.g., cell cycle and proliferation, DNA repair), MYC and PCa-related signatures, stemness, and metabolism (e.g., glycolysis and pyrimidine metabolism) were also, likely, positively regulated by VARS1 (Fig. S2E and S2F). Altogether, these results proposed VARS1 as a proto-oncogenic factor associated with multiple cancer-related pathways to eventually promote PCa progression (also see below for consistent multi-omics dissection and validation).

### 3. VARS1 is essential for PCa maintenance and optimal translation

To experimentally validate the oncogenic role of VARS1, we performed gain- and loss-of-function studies in multiple AR^+^ and AR^—^ PCa cell lines. Lentiviral-mediated *VARS1* knockdown (KD) by short hairpin RNAs (shRNAs) effectively reduced its protein level and the colony formation capability in 4 PCa lines studied (Fig. 2G), with noticeable difference in inhibitory effects among these lines (LNCaP < 22RV1 < DU145 < PC3). Aggressiveness-associated features, such as sphere formation (measuring stemness; Fig. 2H) and migration (Fig. 2I), were also substantially repressed in PCa cells depleting *VARS1*. Again, AR^—^ lines, especially PC3, were generally more sensitive to *VARS1* loss than AR^+^ lines. These results aligned well with the aggressiveness of these 4 lines and the fact that VARS1 is further elevated in CRPC vs. pri-PCa (Fig. 2B and 2C). LNCaP is a relatively indolent and androgen-sensitive pri-PCa-like line [24], whereas 22RV1 is considered as an AR^+^ CRPC [25]. DU145 and PC3 are intrinsically AR^—^ CRPC, with PC3 being the most aggressive line among these four [26]. Consistently, genetic targeting *VARS1* via small interfering RNAs (siRNAs) yielded similar results. Although transient *VARS1* KD by three different siRNAs moderately reduced its protein expression, it obviously inhibited the clonal capacity (Fig. S2G), sphere formation (Fig. S2H), and migration (Fig. S2I) in both AR^+^ and AR^—^ cells, with PC3 and LNCaP being the most and the least affected line, respectively.

Next, we interrogated VARS1’s contribution to protein translation and PCa maintenance in vitro and in vivo. Puromycin incorporation assay revealed that targeting *VARS1* by siRNAs reduced the global translation rates in PCa cells (Fig. 2J). Notably, shRNA-derived stable cell lines via puromycin selection were not suitable for this assay as they were puromycin-resistant. Alternatively, shRNA-mediated *VARS1* KD also diminished total protein biosynthesis measured by polysome profiling (Fig. 2K). Importantly, a reduction in VRAS1 expression (Fig. 2G) also caused a marked decrease in charged proportion of valine-decoding, but not unrelated tryptophan-decoding, tRNAs (Fig. S2J), indicating that the aminoacylation activity is essential for optimal VARS1 function in translation. In vivo tumour regeneration experiments indicated that stable *VARS1* KD inhibited not only the tumour incidence in aggressive AR^—^ CRPC, PC3 (Fig. 2L) and DU145 (Fig. 2M), xenografts but also the tumour growth (growth curve and weight) in both AR^—^ and AR^+^, C4-2 (Fig. 2N) and 22Rv1 (Fig. S2K), models. Obviously, and consistent with the in vitro findings (Fig. 2G), the inhibitory impact of VARS1 deficiency on AR^—^ CRPC tumours was much stronger than that on AR^+^ xenografts, with PC3 being the most sensitive one (Fig. 2L-2N). To further understand what biological processes might be impacted by VARS1 transcriptionally, we performed RNA-seq analysis on PC3 cells with *VARS1* KD (Fig. S2L). Reducing *VARS1* expression by about 3-fold caused misexpression of a total of 837 differentially expressed genes (DEGs; Table S4). GSEA of downregulated genes upon *VARS1* depletion revealed significant enrichment of gene signatures related to cell cycle, DNA repair, tRNA aminoacylation, stemness, and cancer-promoting signaling, among others (Fig. S2L and Table S4), a pattern almost same to *VARS1* gene coexpression analysis as shown in Fig. 2F.

### 4. Aminoacylation activity is required for optimal VARS1 function

To determine whether VARS1’s function in promoting PCa progression was dependent on its tRNA aminoacylation activity, we established stable cell lines expressing wild-type (WT) VARS1 or its catalytic-dead mutant (aa862-865, KMSKS→AMSAS; Fig. 3A). A lack of aminoacylation activity of this mutant (MU) has been previously described [27]. Expectedly, a series of cell-based assays in three different PCa cell lines showed that overexpression (OE) of the WT VARS1 resulted in a significant increase in cell growth (measured by colony formation; Fig. 3B), stemness (by sphere formation assay; Fig. 3C and 3D), and migration (by Trans-well assay; Fig. 3E and 3F) of all three PCa lines. Importantly, MU-OE also increased these aggressive features but to a much lesser extent (Fig. 3B-3F). Tumour regeneration experiments confirmed that VARS1 WT-OE dramatically accelerated tumour growth *in vivo* in both AR^—^ PC3 (Fig. 3G) and AR^+^ 22Rv1 (Fig. 3H) PCa models, with MU-OE displayed intermediate effects. These data highlighted, interestingly, the existence of non-canonical roles of VARS1. In support, such speculation is well aligned with the fact that aaRSs often possess functions beyond tRNA-assisted translation [28] and generally have the ability to catalyze the formation of lysine aminoacylation on its specific substrate proteins, not tRNA, to exert diverse functions [29]. For example, alanyl-tRNA synthetase 1/2 (AARS1/2) can function as lactyltransferase to directly catalyze protein lactylation using lactate and ATP to promote gastric cancer growth via YAP signaling [30] and regulate innate immunity via cGAS [31]. Future studies are needed to look into the possible moonlighting functions of VARS1 (which may not solely rely on aminoacylation activity) in PCa. However, the translational side of VARS1 definitely requires its tRNA aminoacylation activity. Indeed, OE of the WT, but not the MU, VARS1 obviously enhanced the global translation in both AR^+^ and AR^—^ PCa cells as measured by puromycin incorporation assay (Fig. 3I) and polysome profiling (Fig. 3J).

**Fig. 3.**
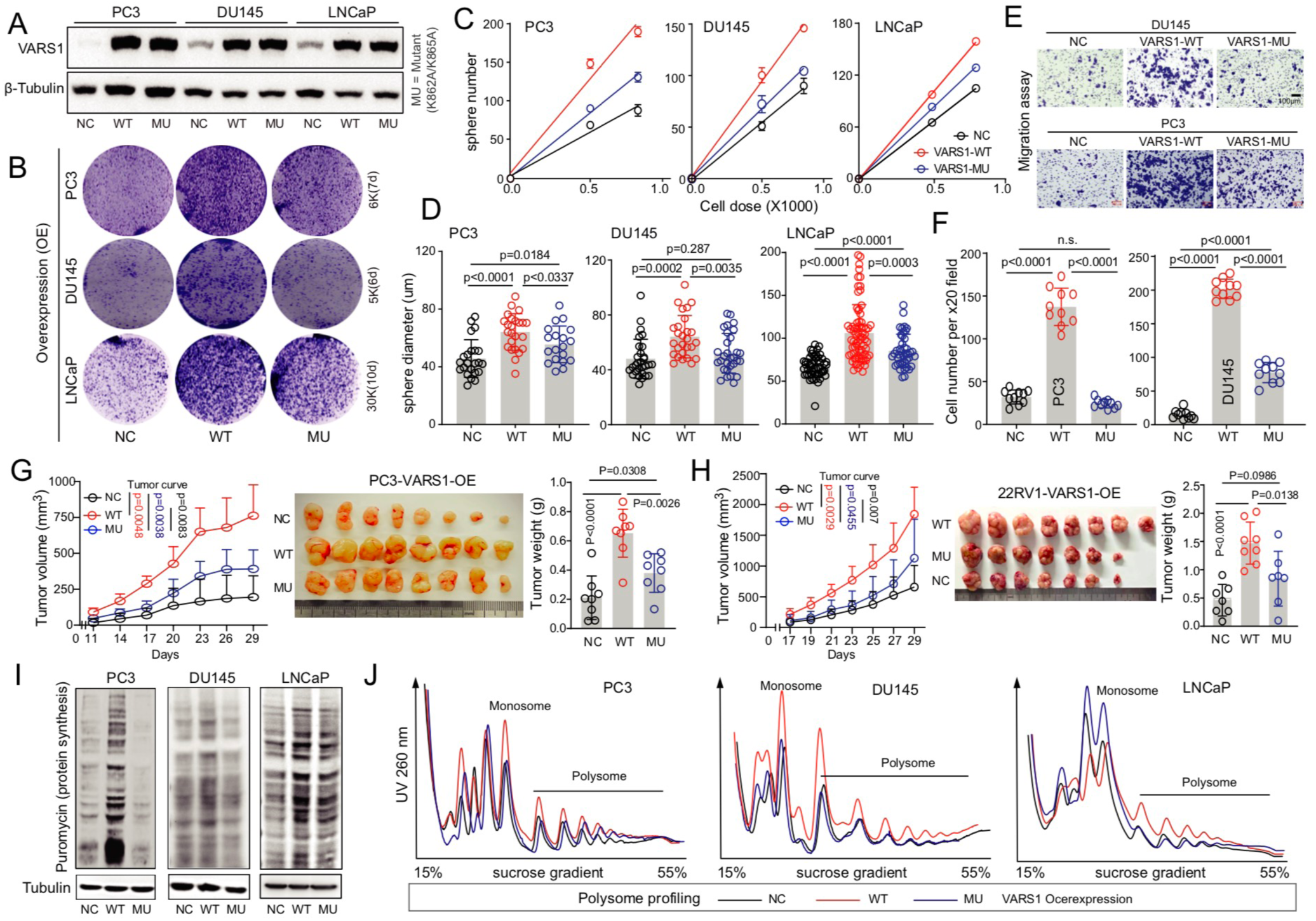
Aminoacylation activity is required for VARS1’s oncogenic function. **A.** Western blot analysis showing overexpression (OE) of WT or catalytically dead (i.e., mutant) VARS1 in indicated PCa cells. β-Tubulin served as a loading control. **B-F**. OE of WT VARS1 significantly, while OE of the mutant form weakly, promotes PCa clonal development (**B**), sphere forming frequency (**C**) and size (**D**), and Trans-well migration (**E** and **F**) in a panel of PCa cells in vitro. For cell migration assay, representative low magnification images (**E**) and quantifications (**F**) are shown. Data represent the means ± SD from cell number counting of 10 random high magnification (20X) images. **G** and **H**. The tRNA aminoacylation activity is required for VARS1 in enhancing PCa development in vivo. Shown are the tumour growth curves (left), endpoint tumour images (middle) and tumour weight (right) of PC3 (n = 8 for each group; **G**) and 22RV1 (n = 7 for control and mutant OE groups, and n = 8 for WT OE group; **H**) models overexpressing mock, WT, or mutant VARS1. Data represent mean ± SD, and the *P*-values for tumour growth curve and weight were determined using two-tailed unpaired Student’s t-test. **I** and **J**. Puromycin incorporation assay (I) and Polysome profiling (J) showing an enhanced global translation in indicated PCa cells with VARS1 WT, but not the mutant, OE. Peaks corresponding to monosome and polysome are labeled.

### 5. VARS1 fuels PCa evolution through codon-selective translational rewiring

The aaRSs ‘charge’ tRNAs by covalently ligating each amino acid to cognate isoacceptor tRNAs [32]. In human, four valine codons (GTA, GTT, GTC, GTG) are decoded by three isoaccepting tRNAs (tRNA-Val^TAC^, tRNA-Val^AAC^, and tRNA-Val^CAC^) respectively, in which both GTC and GTG are decoded by tRNA-Val^CAC^ through wobble pairing. To further dissect the impact of VARS1 on PCa-specific translation dynamics, we utilized multiple translation profiling approaches (Fig. 4A). ***First***, we performed polysome-seq in PC3 cells with or without *VARS1* KD (Fig. S3A and Table S5). Unexpectedly, although depleting *VARS1* reduced global translation (Fig. 2K), it did not specifically, and severely, impact on valine-rich transcripts as we failed to observed a significant enrichment of total valine-codon content in downregulated mRNAs bound by polyribosomes (*p* > 0.05; Fig. 4B). Examination of each valine codon revealed that PCa cells with low *VARS1* preferred to use GTC and GTG, instead of GTA and GTT, codons (Fig. 4B and Fig. S3B). These data highlighted that VARS1 promotes translation with a biased usage of GTA and GTT codons in PCa cells. ***Next***, we employed Ribo-seq to footprint ribosome-protected mRNA fragments (RPFs) to monitor active translation (Table S6). Evaluation of the RPF length distribution and 3-nt periodicity across biological replicates confirmed the high quality of our data (Fig. S3C). Interestingly, the total valine codon content in active translating mRNAs was also unaltered (*p* = 0.22; Fig. 4C). But the codon usage of GTC and GTG, but not GTA and GTT, was markedly increased in upregulated transcripts upon *VARS1* KD in PC3 cells (Fig. 4C and Fig. S3D). ***Last***, we carried out the ultimate proteomics analysis on control and VARS1-depletd cells, whose VARS1 protein levels were reduced by 50% (Fig. S3E and Table S7). Principal component analysis (PCA) showed that the control and *VARS1* KD samples were grouped together and well-separated, leading to identification of 1,590 differentially expressed proteins (DEPs) at a threshold of FDR < 0.05 and FC ≥1.2 (Fig. S3E). Analysis of valine codons indicated that valine-rich proteins were not enriched in downregulated DEPs compared with upregulated ones (Fig. 4D), indicating, again, that targeting VARS1 did not specifically affects translation of valine-rich proteins. Consistently, detailed analysis of four valine codons confirmed a significantly reduced usage of GTA and GTT in VARS1 depleting cells (Fig. S3F). Alternatively, based on aforementioned three distinct omics data, we binned differentially expressed transcripts or DEPs into four quartiles according to the indicated valine codon content, and plotted the log_2_(FC) as a cumulative distribution function. Transcripts encoding genes in polysome-seq (Fig. S3G) and in Ribo-seq (Fig. S3H) and proteins in proteomics (Fig. S3I) with the most GTC/GTG codons were markedly enriched in mRNAs with the top 25% (vs. bottom 25%) codon usage in PC3 cells upon *VARS1* KD, indicating a codon-biased rewiring in valine-rich translation upon VARS1 targeting.

**Fig. 4.**
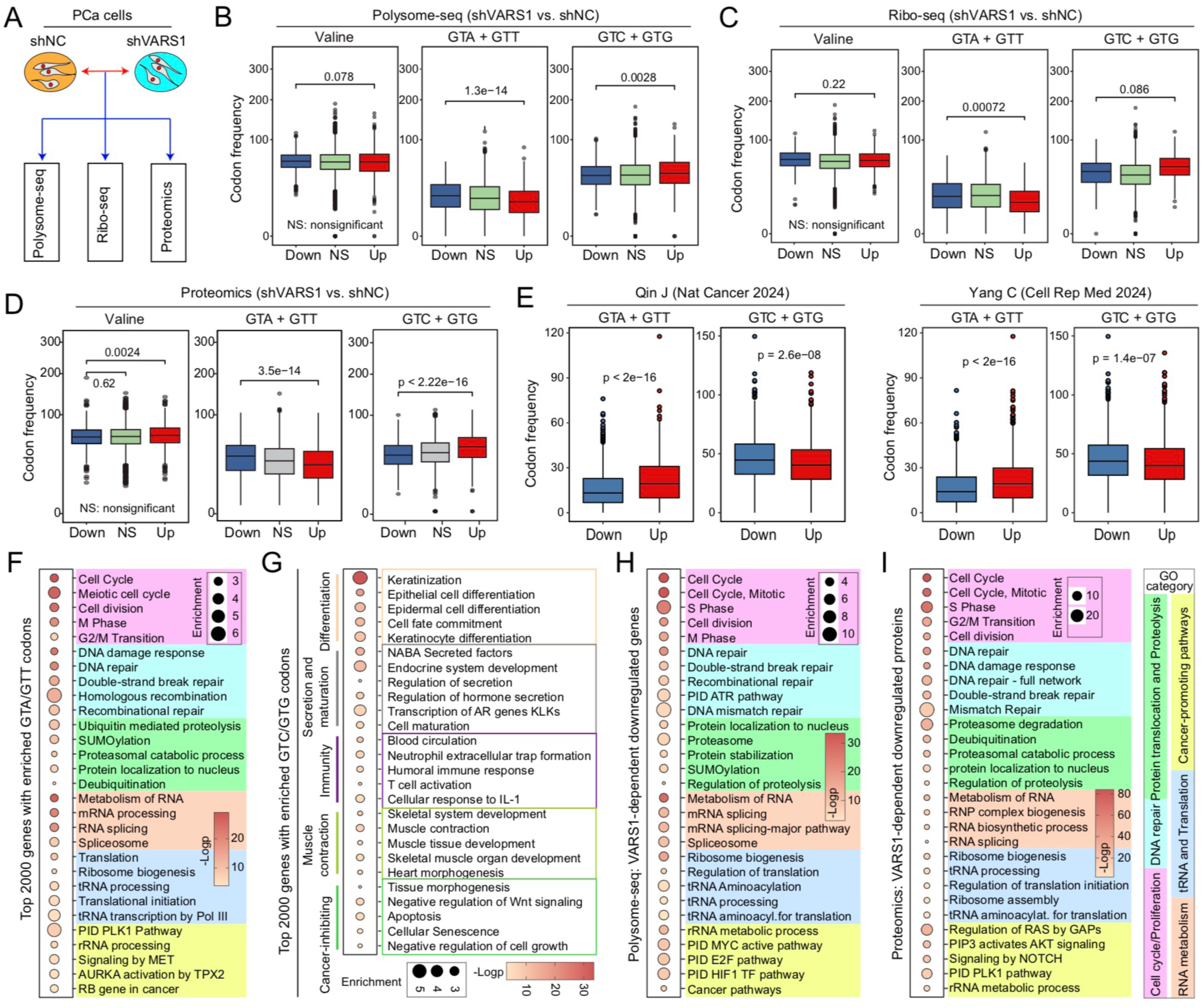
VARS1 promotes PCa through codon-selective translational rewiring. **A.** Schema of experimental approaches to dissect VARS1’s function in translation. **B-D**. Comparison of valine codon usage in DEGs identified in VARS1-depleted vs. control PCa cells by polysome-seq (**B**), Ribo-seq (**C**) and proteomics (**D**). Analysis was stratified by total valine (all four codons combined; left), Val-GUA and Val-GUU (middle), and Val-GUC and Val-GUG (right) codon content, respectively. **E**. Pairwise comparison of valine codon usage in up- and down-regulated proteins identified in indicated proteome cohorts during PCa development (cancer vs. normal). Within the plots, the center lines represent median values, box edges are 75^th^ and 25^th^ percentiles, and whiskers denote the maximum and minimum values, respectively. Significance was calculated by the Wilcoxon test (NS, not significant). **F** and **G**. GO analysis of genes with the most GTA/GTT codons (top 2000; **F**) and GTC/GTG codons (top 2000; **G**) in the human transcriptome. **H** and **I**. GO analysis of VARS1-dependent downregulated genes in polyspme-seq data (**H**) and downregulated proteins in proteomic data (**I**). Terms with p-value <0.01, minimum count 3, and enrichment factor >1.5 (the ratio between observed count and the count expected by chance) were considered significant and all terms were significant. The terms with similar descriptions and functions were grouped into different functional categories, as indicated.

To establish the clinical relevance of VARS1-dependent codon usage rewiring (from GTC/GTG to GTA/GTT), we analyzed the distribution of valine codons in the human transcriptome, finding that GTC/GTG are the ‘optimal’ codons in the normal human transcriptome (Fig. S3J), consistent with a previous report [33]. Cancer usually disrupts the gene expression pattern of healthy tissue resulting in altered codon usage preferences [34]. To understand whether upregulated VARS1 in PCa tissues creates a valine codon-usage ‘switch’, we examined the valine codon contents in DEPs identified in two recently reported large PCa proteome cohorts [10, 12]. Surprisingly, upregulated proteins in clinical specimens utilized GTA/GTT codons more frequently in both cohorts, with GTC/GTG being ‘optimal’ in normal/benign prostates as they were enriched in downregulated proteins (Fig. 4E). Detailed analysis of individual valine codons further supported this notion (Fig. S3K). These results indicated a switch in valine codon-biased usage during prostate tumourigenesis, which is expected to contribute to proteomic reprogramming in PCa. Functionally, Gene ontology (GO) analysis (http://metascape.org) of human genes with the most GTA/GTT codons exhibited an obvious enrichment of many cancer-associated categories (Fig. 4F). The most prominent functional categories were cell cycle/proliferation and DNA damage repair, followed by protein localization and proteolysis, RNA metabolism (mainly splicing), tRNA and translation, and many known cancer-promoting pathways (Fig. 4F). In contrast, genes with the most GTC/GTG content were enriched for pathways tied to differentiation, tissue maturation, and immunity; which were usually associated with cancer-inhibiting functions (Fig. 4G). For instance, we have shown recently that ‘muscle contraction’ category was only and specifically enriched in normal prostates compared to PCa tissues [35]. Furthermore, GO enrichment analysis based on the VARS1-dependent downregulated genes in translation in both polysome-seq (Fig. 4H) and proteomics data (Fig. 4I) identified almost the same functional categories seen in genes with the most GTA/GTT codons (Fig. 4F), consistently establishing GTA/GTT as protumour valine codons that fuel translation of oncogenic transcripts in PCa. Coincidentally, proteins differentially overexpressed in PCa (vs. N) tissues in a large proteomics cohort [10] were also enriched in such an array of functional pathways (Fig. S3L) as seen in Fig. 4F, particularly the tRNA and translation, and DNA repair categories.

### 6. Identification of key VARS1 translational targets in PCa

To identify candidate genes that mediate VARS1’s cancer-promoting functions, we overlapped the VARS1-dependent downregulated genes and proteins identified in polysome-seq and proteomics data, respectively, with genes carrying the most GTA/GTT codons (Fig. 5A), yielding seven candidates. Among them, ubiquitin conjugating enzyme E2 T (UBE2T), a subunit of fanconi anemia (FA) core complex [36], and lin-9 DREAM MuvB core complex component (LIN9), a putative tumour suppressor that inhibits DNA synthesis [37], have been known to participate in DNA repair pathway. The kinesin family member 20B (KIF20B) [38] and structural maintenance of chromosomes 4 (SMC4) [39] are involved in cell-cycle and proliferation regulation. As cell cycle/proliferation and DNA damage repair were the categories repeatedly present in VARS1 positively regulated genes (Fig. 4F, 4H, 4I), we thus focused on these two oncogenic categories. In the TCGA cohort [7], expression of LIN9 and SMC4 was not significantly and consistently associated with PCa pathological grade and patient survival (Fig. S4A). However, expression of UBE2T and KIF20B was positively and negatively associated with the tumour grade and overall survival outcome, respectively (Fig. S4B), indicating potential oncogenic roles in PCa. We therefore chose UBE2T and KIF20B for further characterization. Western blot analysis confirmed reduced UBE2T and KIF20B protein expression in VARS1 KD cells, but without affecting their mRNA levels (Fig. 5B). Moreover, we performed quantitative reverse transcriptase PCR (RT-qPCR) on the mRNAs extracted from sucrose gradient fractions, and revealed that the abundance of UBE2T and KIF20B mRNAs was decreased in the heavy polysome (actively translating) fractions in PC3 cells upon VARS1 depletion; whereas the mRNAs of housekeeping genes (i.e., GAPDH and α-Tubulin) showed no changes in their distribution pattern (Fig. S4C). Together, these data suggested that downregulation of these proteins is mainly due to VARS1-mediated translational alterations.

**Fig. 5.**
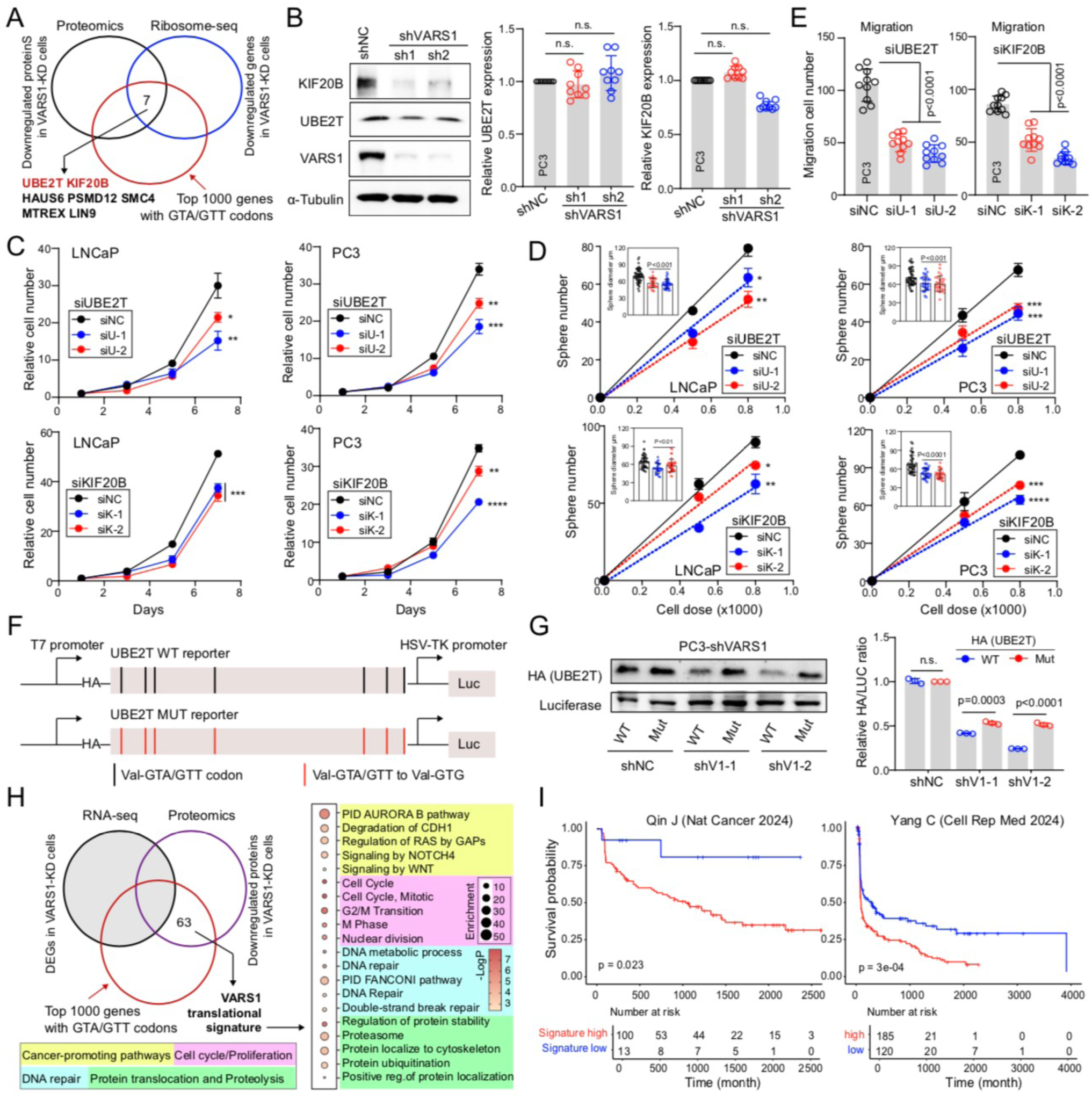
VARS1 regulates translation of oncogenic transcripts in a codon-biased manner. **A**. Three-way overlap between indicated gene sets. **B**. Western blot (left) and qPCR (right) analysis showing that depleting VARS1 reduces the expression of KIF20B and UBE2T at the protein, but not mRNA, levels. **C-E**. Small interfering RNAs (siRNAs)-mediated knockdown of indicated genes severely inhibits proliferation (15k/well for PC3 and 30k/well for LNCaP were plated in 12-well plates with four repeats for each condition, respectively; **C**), sphere formation (**D**) and migration (**E**) in both AR^+^ LNCaP and AR^—^ PC3 cells. Two individual siRNAs for each gene were used. **F**. Schematic overview of UBE2T codon reporter design. The coding sequence (CDS) of this gene is short (504nt). All PCa-preferred valine GTA/GTT codons were replaced with GTG codons. WT, wild type; MUT, mutant. **G**. Western blot for UBE2T reporter expression (HA expression) in PC3 VARS1-depleted cell lines. Luciferase is used as a transfection control. Shown are representative of n = 3 independent experiments (left) and quantification data (right). Statistics calculated by unpaired two-tailed Student’s t test. **H**. Identification of a VARS1 translational signature. Shown are three-way overlap between indicated gene sets (left) and GO analysis of the VARS1 translational signature (right). **I**. A higher level of VARS1 translational signature correlating with reduced overall patient survival in two PCa proteome cohorts.

To determine whether these two proteins indeed play oncogenic roles in PCa, we used siRNA-mediated depletion technology. KD of *UBE2T* or *KIF20B* in both AR^—^ PC3 and AR^+^ LNCaP cells (Fig. S4D) inhibited cell viability (Fig. 5C), clonal capacity (Fig. S4E), and sphere formation (both sphere number and size (inserted); Fig. 5D). Also, PC3 cells showed a decreased migration ability upon VARS1 depletion (Fig. 5E and Fig. S4F). These findings establish UBE2T and KIF20B as VARS1-regulated tumour promoters in PCa. Finally, to determine if VARS1 overexpression in PCa can mediate codon-biased translation of a downstream target gene, we performed valine codon-based mutagenesis studies of UBE2T as its coding sequence (CDS) is short (only 504nt). The reporter constructs containing UBE2T were designed where the GTA/GTT were mutated to the synonymous codon GTG, a codon for which translation was preferred upon VARS1 KD (Fig. 5F). We observed that, while the translation of WT transcript was markedly reduced, the codon mutant form of UBE2T was less repressed, by VARS1 depletion (Fig. 5G), consistent with direct VARS1-dependent and valine-codon-biased regulation of UBE2T. Collectively, these results confirmed, again, that VARS1 KD reverses the valine codon-dependent translation from a preferred GTA/GTT usage in PCa to a selective GTC/GTG usage which are preferred in normal prostates.

Finally, to further relate the VARS1-dependent translational program observed in PCa cells to clinical relevance, we performed RNA-seq analysis on VARS1-depleted PC3 cells (Table S4), and integrated the proteomics data and transcriptomics data. We defined the VARS1-translational signature (*n* = 63; Table S8) as genes (1) downregulated at protein levels, (2) whose mRNA expression was unaffected by VARS1 KD (non-DEGs) and (3) enriched in GTA/GTT codons (top 1000 genes used) (Fig. 5H, left). Consistently, the VARS1 signature was predominantly enriched in pathways associated with cancer-promoting signaling, cell cycle/proliferation, DNA repair and proteolysis (Fig. 5H, right). Importantly, such VARS1 signature was predictive of worse patient outcomes in two large PCa proteome cohorts (Fig. 5I), further confirming the protumour role of VARS1-mediated translational rewiring.

### 7. Dietary VR inhibits tumour growth and metastasis

Given the fact that inhibitors directly targeting VARS1 are not currently available and valine is an essential amino acid that must be absorbed through diet, we therefore asked whether valine restriction (VR) could be an alternative anti-cancer strategy. Supplementation of *L*-valine in the culture medium, in which both FBS and basal medium completely lacks valine, increased PCa cell proliferation in a dose-dependent manner (Fig. S5A). Notably, PC3 grew slowly even at increasing valine (Fig. S5A and S5B). Expectedly, no PCa cells survived in the medium when valine was deprived completely (Fig. S5B). Next, we examined the safety and efficacy of dietary VR in vivo. Consistently, prolonged total valine deprivation resulted in life-threatening weight loss in mice (Fig. S5C). According to the minimal requirement of dietary valine for relatively normal mice physiology [14], we tested two intermediate doses of valine by feeding mice a control diet (8g/kg) or a valine-deficient diet supplemented with 0.8g/L (0.08%) or 0.45 g/L (0.04%) valine in drinking water. We did not observe significant weight loss in either immune-deficient nude mice (Fig. S5D) or immune-intact C57BL/6 mice (Fig. S5E) during the experimental period, indicating the safety of VR treatment. As schematically illustrated in Fig. 6A, treatment of castration-resistant PC3 xenografts (Fig. 6B) and RM1 allografts (Fig. 6C) with VR effectively inhibited tumour growth, with two intermediate doses displaying similar inhibitory effects. As MYC overexpression represents a critical driver of PCa progression [24], we also treated the Myc-driven spontaneous murine PCa (Hi-Myc tumours) with VR (0.8g/L) and observed significant inhibition of Hi-Myc tumour growth in vivo (Fig. 6D). Histological examination of whole-mount prostate images revealed large areas of advanced adenocarcinomas in mice fed the control diet; whereas VR-treated Hi-Myc prostates exhibited reduced tumour areas and prominent benign and hyperplasic glands (Fig. 6E and S5F). Importantly, the VR diet did not cause body weight loss or noticeable abnormalities in any major organs examined including heart, liver, spleen, lung and kidney (Fig. S5G). Using cancer cells tail vein injection as a relevant lung metastasis model, we determined the possibility of dietary VR as an anti-metastasis therapeutic regimen. Strikingly, our results showed that VR at slightly less restriction levels (0.8g/L or 0.45 g/L) substantially decreased metastatic burden in lungs compared to mice fed the normal diet, again without animal weight loss (Fig. 6F and S5H). Overall, these data provide compelling evidence that controlled dietary VR can function as an intervention to halt PCa progression without the adverse effects of total valine deprivation.

**Fig. 6.**
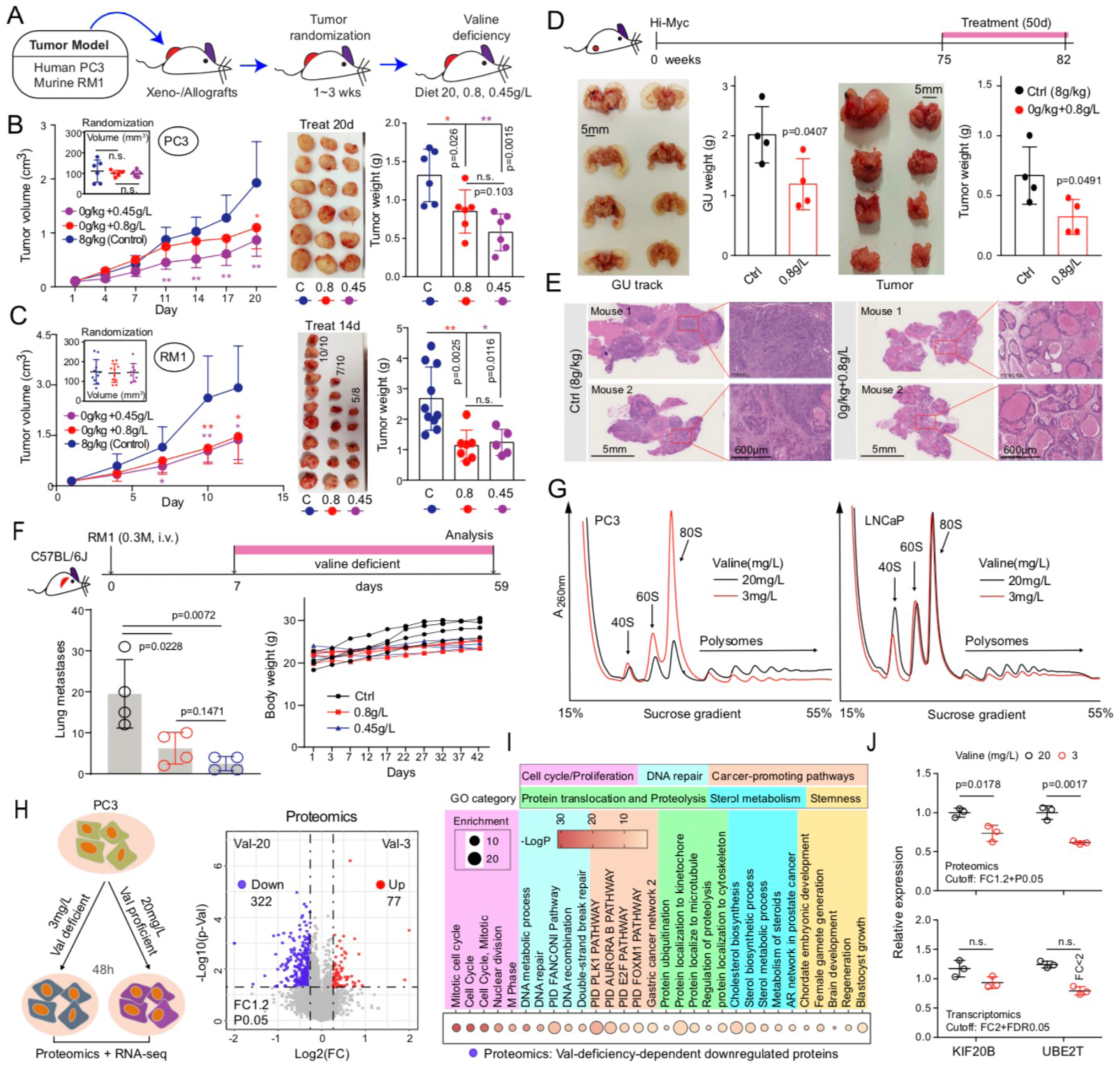
Dietary VR inhibits tumour growth and metastasis. **A.** Schematic of in vivo VR treatment. **B** and **C**. Inhibitory effects of VR on the growth of indicated CRPC models in vivo. Shown are the tumour growth curves (left; insets present tumour randomizations), endpoint tumour images (middle), and tumour weight (right) of PC3 (n = 6 for each group) and RM1 (n= 10 and 8 for vehicle and 08.g/L group, and 0.45g/L group, respectively) models treated with vehicle or VR. Data represent mean ± SD and all *P*-values were determined by two-tailed unpaired Student’s t-test. *P < 0.05. **P < 0.01. ***P < 0.0001. **D.** Schematic of in vivo VR treatment (0.8g/L) in the Hi-Myc PCa model. Gross images and weight of the genitourinary (GU) tract (left) or the prostate (right) in Hi-Myc mice treated with vehicle or VR. **E.** Representative whole-mount images of hematoxylin and eosin-stained prostate sections from two Hi-Myc mice treated with vehicle or VR, respectively. Boxed regions are enlarged. **F.** Dietary VR inhibits PCa metastasis. A tail-vein injection model of lung metastasis is used for the VR treatment design (top). VR, compared with vehicle, reduces the murine RM1-induced lung metastasis burden but not the body weight of indicated models treated with vehicle or VR (bottom). *P*-values were determined by two-tailed unpaired Student’s *t*-test. **G.** Polysome profiling in PCa cells cultured in low (3 mg/L) and high (20 mg/L) valine medium showing impact of in vitro VR on translation. Peaks corresponding to 40S small subunit (40S), 60S large subunit (60S), monosome (80S), and polysome are labeled. **H.** Multi-omics analysis of PC3 cells cultured in control or VR medium for 48 h. Shown are experimental schematic (left) and effect of VR on PCa proteome in vitro (*n* = 3 per group; right). **I.** GO analysis of valine deficiency-dependent downregulated proteins in proteomic data. **J.** Relative expression of KIF20B and UBE2T at both protein (top) and mRNA (bottom) levels as revealed by proteomics and RNA-seq analysis in PC3 cells cultured with or without valine depletion, respectively. Statistics calculated by unpaired two-tailed Student’s t test.

Next, we examined whether VR exerts anti-PCa function through, at least partially, translational rewiring. PCa cells cultured in a low valine medium (3mg/L for 2-3 days, a concentration that significantly inhibited proliferation) exhibited a marked decrease in translation rates compared to them maintained in a high valine medium (20mg/L), as revealed by polysome profiling (Fig. 6G). Notably, the translation of AR^—^ PC3 cells were affected more severely than that in relatively indolent AR^+^ LNCaP cells. Proteome profiling revealed that a total of 399 DEPs were identified in valine-deficient cultures (Fig. 6H and Table S9), with majority (81%) being downregulated. Importantly, downregulated proteins were predominantly enriched for pathways tied to cell cycle/proliferation and DNA repair (Fig. 6I), followed by cancer-promoting category and protein localization and proteolysis. Besides, categories of stemness and steroid hormone metabolism were also enriched in such downregulated proteins, indicating reduced stemness along with and enhanced differentiation of AR^—^ PC3 cells toward an androgen-sensitive state upon valine deficiency, respectively (Fig. 6I). Interestingly, valine deficiency (3 vs. 20 mg/L) had little impact on global mRNA expression, as we only observed 23 DEGs at a stringent statistical threshold of ≥2 FC and FDR of <0.05 (Table S10). The expression of two VARS1 translational targets were reduced at protein, but not at mRNA, levels (Fig. 6J), again supporting that limited valine bioavailability decreases translation via VARS1.

### 8. VARS1 inhibitor suppresses growth of aggressive PCa

To investigate the therapeutic potential of VARS1 inhibition in aggressive PCa, we attempted to develop a small molecular inhibitor of VARS1 by performing a virtual high throughput-screening using a molecular docking model [40] with an AlphaFold-2 predicted three-dimensional (3D) structure of the VARS1 (UniProt ID: P26640; Fig. 7A), as no experimentally solved structure has been reported yet. We first screened approximately 10^6^ small molecule probes from the Specs chemical library and chose the top 100 hit compounds that potentially bound VARS1 for further analysis (Table S11). After removing structurally similar compounds, the structural diverse selection resulted in 42 chemicals left, of which 35 were purchasable (Table S11). To initially screen the cell-killing effect of these compounds, we performed MTT assay and found that, when PCa cells were treated at 10 µM concentration, 18, 20, and 24 out of 35 compounds were toxic to PC3, DU145, and LNCaP cells, respectively, at a threshold of cell growth inhibition ≥20% (Fig. S6A). Overlap of these effective hits revealed 13 compounds as potential anti-PCa agents. Next, we performed fluorescence quenching assay to test whether these 13 compounds actually bind, and preferentially bind, to bacterially purified VARS1 over its paralog VASR2 protein. Consequently, 6 out of 13 strongly bond to VARS1 (Fig. S6A and S6B), but only 3 specifically bond to VARS1 as some compounds displayed similar affinity to both purified VARS proteins in vitro (Fig. S6C). Structural prediction of VARS1 binding pockets to the 3 candidates indicated that both compound 2701 and 2901 can interact with VARS1 through hydrogen bonds at residue D824, P343 and N345, respectively, with no obvious and specific interaction observed for 3201 (Fig. S6D). Notably, these three interacting residues were all located in the aminoacylation domain (Fig. S6D). Colony formation assay highlighted that 3201 did not exhibit strong anti-proliferation effects even at relatively high concentrations such as 15 µM (Fig. S6E). Therefore, we next focused on 2701 and 2901 as potential VARS1 inhibitors (VARS1i) for further characterization.

**Fig. 7.**
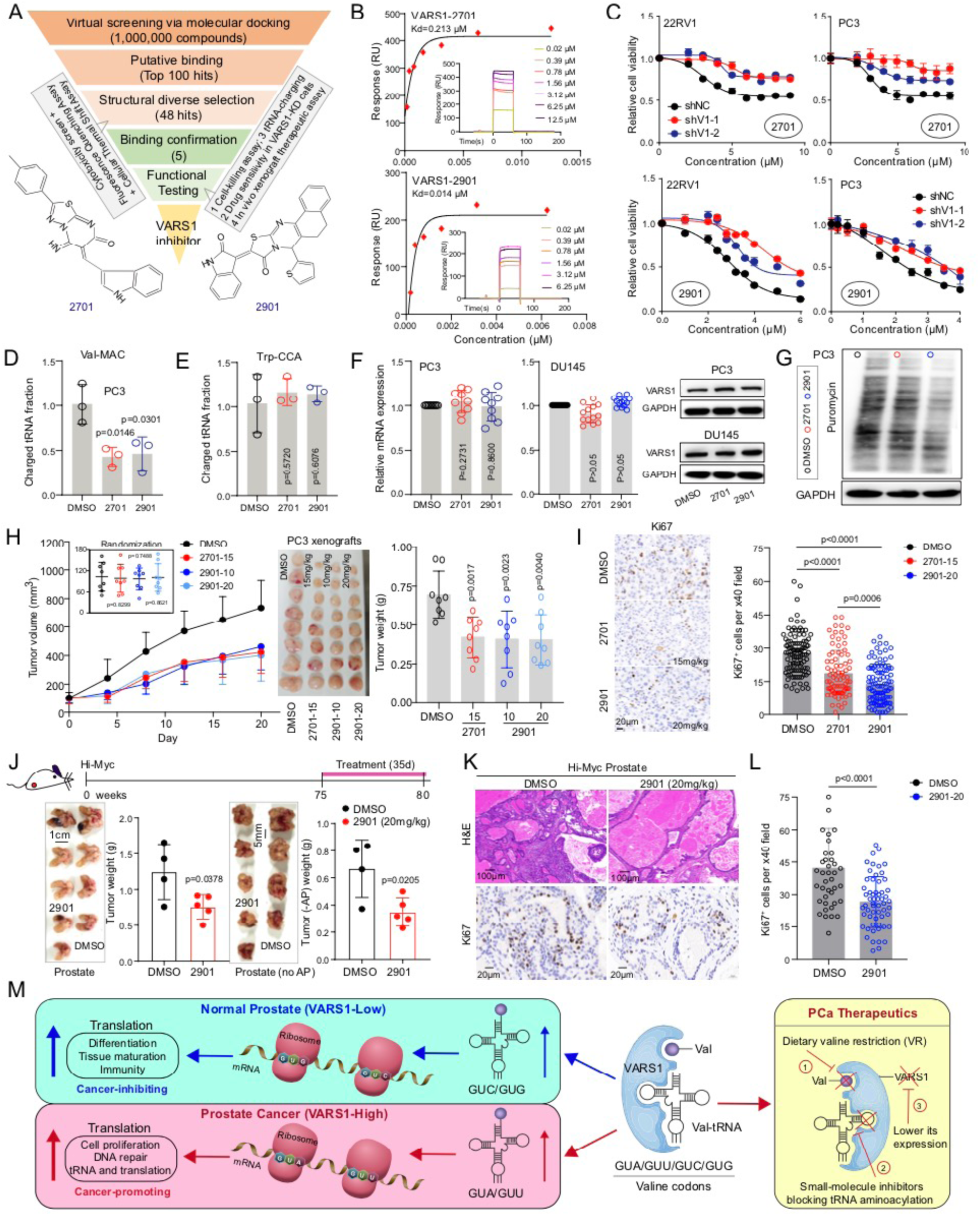
VARS1 inhibitor suppresses tumour growth. **A.** Scheme of virtual screening protocol for small-molecule inhibitors of VARS1. The chemical structure of 2701 and 2901 are shown. **B.** SPR equilibrium binding of the indicated small molecules to bacterially purified VARS1. Insets present SPR sensorgrams for indicated molecules binding to VARS1 protein (with fit to 1:1 model). **C.** Cell viability assay showing growth inhibition of AR^+^ 22RV1 (2k/well) and AR^—^ PC3 (1.5k/well) cell lines with or without VARS1 depletion after 3 days treatment with serial dilutions of indicated molecules. **D** and **E**. Treatment of both 2701 (10 μM for 2 days) and 2901 (2 μM for 2 days) reduces the charged to total ratio of Val-tRNA^MAC^ (M=A or C; D), but not the ratio of irrelevant Trp-tRNA^CCA^ (E). **F.** Expression of VARS1 at mRNA (left) and protein (right) levels in indicated cells treated with 2701 (10 μM for 2 days) or 2901 (2 μM for 2 days), as revealed by qPCR and western blot analysis. **G.** Puromycin incorporation assay showing a visible reduction of total protein synthesis rate in PC3 cells upon 2701 (10 μM) or 2901 (2 μM) treatment for 2 days. **H.** Inhibitory effects of VARS1i on the growth of PC3 xenografts. Shown are the tumour growth curves (left; insets present tumour randomizations), endpoint tumour images (middle), and tumour weight (right) of PC3 model treated with vehicle or indicated VARS1i (*n* = 8 per group). **I.** Representative IHC images of Ki67 staining for proliferation (left) and quantification data (right) in indicated endpoint tumours. **J.** Schematic of in vivo 2901 treatment in the Hi-Myc PCa model. Shown are gross images and weight of the genitourinary (GU) tract (left) or the prostate without AP (right) in Hi-Myc mice treated with vehicle or 2901. **K.** Representative H&E (top) and Ki67 (bottom) staining of prostate sections from Hi-Myc mice treated with vehicle or 2901, respectively. **L.** Quantification of Ki67^+^ cells in indicated endpoint tumours. Data represent mean ± SD from cell number counting of >40 random high magnification (x40) images. The P value was calculated using Student’s t-test. **M.** Model of VARS1-mediated acceleration of PCa development and progression, with potential therapeutic approaches indicated.

Surface plasmon resonance (SPR) results again confirmed a strong interaction between the two probes and VARS1, revealing that 2701 and 2901 directly bound to VARS1 with a kinetic *K*_d_ of 0.213 and 0.014 µM, respectively (Fig. 7B). As expected, cell-based assays indicated that both compounds significantly inhibited cell growth at a dose-dependent manner (Fig. S6F), with 2901 displaying a superior effect and AR^—^ CRPC cells (especially PC3) generally being more sensitive to these drugs. To further confirm that VARS1 is indeed a functional target, we treated PCa cells with or without *VARS1* KD with these compounds, finding that the cell killing effects of them were significantly reduced when cells lacking *VARS1* (Fig. 7C). Notably, 2901, relative to 2701, at lower concentrations generated greater inhibitory effect on cell growth. Due to an essentiality of this protein to PCa cell survival, we failed to generate complete *VARS1*-knockout cells. We thus used shRNA-mediated KD instead, which may explain why cells with *VARS1* KD were still moderately responding to these two molecules compared to control cells due to the existence of residual VARS1. Biochemically, we also examined charged tRNA-Val^MAC^ (M=A or C) levels in PCa cells and observed a reduction in charged to total tRNA-Val ratio when treated with these drugs (Fig. 7D). The charging of irrelevant tRNA-Trp^CCA^ was unaffected (Fig. 7E). Interestingly, VARS1 expression either at mRNA or protein level was not altered after 2701 (10 µM for 2 days) or 2901 (2 µM for 2 days) treatment (Fig. 7F). Puromycin incorporation assay consistently revealed that VARS1i suppressed the protein biosynthesis in PCa cells, again, with 2901 being more effective (Fig. 7G). Altogether, these data suggested that 2701 and 2901 exerted anti-PCa effects mainly through interfering VARS1’s tRNA aminoacylation activity but not its expression.

Last, to assess the translational potential of VARSi in vivo, we treated PC3 xenografts in immune-deficient hosts (15 mg/kg for 2701 (note that the available amount for this probe is limited for purchase), 10 and 20 mg/kg for 2901 via intraperitoneal (i.p.) injection for once the other day). Both drugs at indicated doses generated comparable effects in inhibiting tumour growth in vivo in terms of tumour growth, size and weight (Fig. 7H). No overt toxicity of VARS1i to mice was observed, as indicated by stable body weights (Fig. S6G) and sizes of major organs (Fig. S6H) in control and drug treated groups. Immunostaining of Ki-67 indicated a decreased cell proliferation in endpoint tumours treated with VARSi vs. vehicle (Fig. 7I). Furthermore, we also treated the spontaneous Hi-Myc animals with vehicle and 2901 (20 mg/kg) for 35 days and, consistently, observed an obvious inhibition of late-staged tumour growth in vivo (Fig. 7J). Importantly, both the whole prostates and prostates without anterior prostate (AP) lobes were decreased in weight (Fig. 7J). Tumour development in AP is slow due to a low activity of probasin promoter in such lobes [41, 42]. H&E images confirmed that the aggressiveness of Hi-Myc tumours was markedly suppressed histologically after 2901 treatment (Fig. 7K_upper_ and Fig. S7A), together with a decreased Ki-67 proliferation index (Fig. 7K_bottom_ and 7L). Again, VARSi treatment did not change the weight of mouse bodies and many major organs examined (Fig. S7B), indicating drug safety. Collectively, our data established VARS1 as a valid target in advanced PCa, urging the potential development of VARS1i into clinical use.

## DISCUSSION

Here, we identify that VARS1 is continuously overexpressed during human PCa evolution (from development to progression) and correlates with a worse patient survival in clinical data. Loss- and gain-of-function studies places VARS1 as an oncogene to promote translation of GTA/GTT codon-enriched transcripts in PCa; whereas GTC/GTG are the favored ‘optimal’ valine codons in normal or benign prostates (Fig. 4). Functionally, VARS1-biased codon usage elevates translation of valine-rich genes that are strongly implicated in proliferation (such as cell cycle and DNA repair pathways) and many cancer-promoting pathways (Fig. 7M). One such gene, UBE2T, a valine-enriched protein involved in protective DNA repair, is positively regulated by VARS1 at translation level. UBE2T is also overexpressed in human PCa and biologically sustains malignant mitosis and stemness. Of clinical relevance, we also define a VARS1 specific translational signature predictive of adverse clinical features. Depletion of VARS1, or dietary restriction of valine, or small-molecule VARS1i markedly suppresses global translation and dampens PCa growth and metastasis in vivo (Fig. 7M).

PCa displays a distinct mRNA translation programme, leading to an aberrant proteome that fuels tumour development [9–12]. A recent proteomic profiling study revealed that proteins upregulated in pri-PCa samples are enriched in various protumour pathways, such as translation initiation, ribosome biogenesis, amino acid metabolic process, protein localization and proteolysis, and RNA metabolism (including splicing) [11]. Consistently, systematic GO analysis of upregulated proteins (FC ≥ 1.2) in high-risk PCa in another proteome cohort [10] also highlights significant enrichment in similar functional categories linked to cell mitosis, DNA repair, protein localization and proteolysis, RNA metabolism, tRNA and translation, and cancer-promoting pathways (Fig. S3L). Strikingly, almost the same categories are found enriched in genes with enriched GTA/GTT codons, in genes bearing low translation activity (polysome-seq), and in proteins downregulated in PCa cells (proteomics) upon VARS1 depletion (Fig. 4). Importantly, the VARS1 signature is also enriched in these pathways, strongly suggesting that VARS1 is the main, or at least in large part, driver of reprogramming the proteomic landscape during PCa evolution. In particular, the translational targets of VRAS1 impact a spectrum of pathological processes that function convergently to accelerate PCa progression.

We found that although OE of mutant VARS1 causes no noticeable enhancement in protein biosynthesis, it does increase, mildly, PCa fitness both in vitro and in vivo in multiple biological assays (Fig. 3). It is in line with the fact that many aaRS serve cellular functions independent of their roles in aminoacylation and canonical translation [28, 32]. Therefore, the possibility that VARS1 may exacerbate PCa through a non-tRNA dependent mechanism cannot be fully excluded. While this hypothesis remains to be tested, it is unlikely that the translation-independent function of VARS1 plays a dominant role in PCa progression based on our findings. In parallel, it is worth noting that VARS1 may exert context-dependent functions in different pathological settings. It has been reported that valine tRNA biogenesis promotes tumourigenesis and a worse survival in leukaemia. And, *VARS1* KD reduced translation rates of mRNAs that encode subunits of mitochondrial complex I, leading to defective assembly of complex I and tumour growth due to impaired oxidative phosphorylation [14]. However, we did not observe an obvious and consistent enrichment of mitochondrial complex I genes in VARS1-regulated targets at translation level in our system, as revealed by our multi-omics data, suggesting the mechanistic difference in hematological vs. solid cancers. Interestingly, another recent study has linked VARS1 more restrictedly to drug resistance in melanoma. Depleting VARS does not significantly compromise melanoma cellular survival nor tumour growth in mice, unless cells or tumours are treated with MAPK therapy [33]. Mechanistically, VARS1 specifically protects melanoma from targeted therapy via promoting *HADH* translation, a valine-enriched enzyme, which in turn upregulates fatty acid oxidation to promote resistance [33]. Here we show that VARS1 is upregulated in pri-PCa at both mRNA and protein level, and its KD impacts negatively both pri-PCa-like AR^+^ LNCaP and CRPC PC3 cells, albeit at different extent with LNCaP being less affected. Our study and others commonly highlight that VARS1 might play a more prominent role in cancer progression, especially at a late-stage of drug resistance. Interestingly, we demonstrate that *VARS1* KD in PCa decreases metastasis and actually switches the valine codon usage from protumor GTA/CTT codons to normal tissue favored GTC/GTC codons, further suggesting the safety of drugging VARS1 in a cancer setting.

Given the importance of translational regulation, and the fact that many aaRSs are misexpressed, in human cancers [43], studies have been proposed aaRSs as promising therapeutic targets [27, 32]. Notably, aaRSs have been a focus of research for antimicrobial drug discovery in the past decade, with several aaRS inhibitors (aaRSi) being in clinical practice already [44]. For instance, GSK3036656 (ganfeborole), a first-in-class benzoxaborole inhibiting the *Mycobacterium tuberculosis* LARS1, has demonstrated a promising bactericidal activity and acceptable safety profile in a phase IIa clinical trial [45]. However, development and subsequent use of aaRSi for cancer treatment are only just beginning to be understood. Febrifugine derivatives, including Halofuginone which target EPRS and inhibit prolyl-tRNA synthetase activity, are under investigation for the treatment of cancer and inflammatory diseases [46]. Moreover, another catalytic inhibitor of EPRS, NCP26, has recently developed and demonstrated anti-myeloma activity in preclinical models [19]. A previous study, based on an in vitro screening of a library of 1,3-oxazines, benzoxazines and quinoline scaffolds, reported several compounds capable of suppressing proliferation of the lung and colon cancer cells via targeting methionyl-tRNA synthetase (MARS) [47]. Unfortunately, none of these aaRSi have reached appreciable stages of clinical trials against human cancers. Currently, there is no small-molecule VARS1i available to our best knowledge. To bridge this knowledge gap, we developed chemical probe 2901 as a novel inhibitor that targeted the aminoacylation, so as the translational, activity of VARS1, and demonstrated the preclinical efficacy in halting prostate tumour growth and metastasis. In summary, we highlight VARS1 as a ‘translational’ driver of PCa evolution and uncover attractive therapeutic opportunities for patients with aggressive PCa.

## METHODS

### Cell culture and in vitro VR treatment

The RM1, LNCaP, 22RV1, PC3, and DU145 cell lines were obtained from the National Collection of Authenticated Cell Cultures (Shanghai, China). These cell lines were maintained in RPMI-1640 medium (Gibco, 8123209) supplemented with 10% FBS (ExCell Bio, FSP500, China) and 1% penicillin–streptomycin. All cells were maintained at 37°C and 5% CO_2_ in a humidified incubator. For valine deficiency experiments in vitro, *L*-Valine (sigma, V0513) with different concentrations were added to valine-deficient RPMI-1640 culture medium (Rigorous Scientific, China) containing 10% dialyzed FBS (VivaCell biosciences, C3820-0100) and 1% penicillin–streptomycin.

### Cell proliferation, colony formation, cell migration, and sphere assays

For the cell proliferation assay [24], cells were seeded in triplicate into 12-well plates and cell number was quantified every 2–3 days using a hemocytometer. For inhibitor studies, we usually plated cells in normal medium at day 1, and then added the inhibitors at varying concentrations on day 2 to avoid the drug’s interference with cell adhesion. For colony formation assays [24, 48], we generally plated PCa cells at a low density (i.e., 4K cells per well for PC3, 6K cells for DU145, 20K cells for 22RV1, and 30K cells for LNCaP) into 6-well plates and let cells grow for 7–10 days before visualization of the culture by 0.1% crystal violet staining (Sigma, C6158). For cell migration assay, usually 40–50K cells suspended in 200 μL of medium containing 1–2% FBS were loaded onto the upper chamber of Transwell inserts (Corning, 3422). The lower chamber contained culture medium supplemented with 20% FBS as a chemoattractant. After incubation for 16–40 h, the Transwell chambers were subjected to 4% paraformaldehyde fixation and crystal violet staining. For sphere formation assays [2], cells were suspended in 1:1 Matrigel (BD Biosciences)/RPMI-1640 in a total volume of 100 μL. The mixtures were then plated around the rim of wells in a 12-well plate and allowed to solidify in a 37°C incubator for 30 min, followed by the addition of 1 mL of warm SC medium [DMEM/F12 supplemented with insulin (4 μg/mL), B27 (Invitrogen), and epidermal growth factor (20 ng/mL), and basic fibroblast growth factor (10 ng/mL)]. Usually 8–11 days after plating, spheres with a diameter of more than 50 mm were counted.

### Histology and IHC

All human tissue slides from PCa patients were obtained from Zhongda Hospital Southeast University with informed consent from patients. The use of clinical samples is under the supervision of Ethics Committee of Zhongda Hospital, Southeast University (approval ID 2022ZDKYSB099) and in accordance with the committee’s guidelines. H&E and IHC staining were performed on 5-mm paraffin-embedded sections. For IHC staining, slides were deparaffinized in xylene and hydrated in gradient alcohols to water. Endogenous peroxidase activity was blocked with 3% H_2_O_2_ for 10 min, followed by antigen retrieval in 10 mM citrate buffer (pH 6.0). After blocking with Biocare Blocking Reagent (Biocare), slides were incubated with primary antibodies (Table S12), followed by incubation with secondary antibodies and DAB (Bio-Genex Laboratories) development [2]. Data quantification for IHC staining was based on the cell number counting of many random high magnification images (as indicated in the figure legends) derived from 3 to 5 tumors per condition.

### Western blotting, RNA isolation and quantitative RT-PCR

Standard western blotting procedure was used. Briefly, Cells were lysed and total protein concentrations in supernatants were quantified using Bicinchoninic Acid Assay (BCA) kit (Beyotime Biotechnology, Shanghai, China). The proteins were denatured and fractionated by SDS-PAGE, then transferred onto PVDF membrane (Millipore, Billerica, USA), blocked in 5% nonfat milk, and incubated with specific primary antibody (Table S12) at 4°C overnight. After washing with PBST, the membrane was incubated with secondary antibody, exposed by Hyperfilm ECL kit and finally images were acquired. Total RNA was extracted using the RNA-easy Isolation Reagent (Vazyme, R701) following the manufacturer’s guidelines. cDNA was synthesized using the HiScriptII Q RT SuperMix for qPCR (+gDNA wiper) (Vazyme, R223). cDNA was then used for quantitative PCR by Genious 2X SYBR Green Fast qPCR Mix (ABclonal, RK21204) on QuantStudio 1 (Thermo Fisher Scientific, America) system. All primers used in the present study were provided in Table S12.

### siRNA, shRNA, and overexpressing plasmids

The siRNAs targeting human VARS1 were designed and synthesized by Sangon Biotech (Shanghai, China). siRNAs were then introduced into cells at a concentration of 80 nM using Lipofectamine RNAi MAX in an Opti-MEM medium and cultured for >36 h before used for downstream experiments. For shRNA-mediated KD experiments, we used pLKO.1-EGFP-Puro lentiviral vector to construct the VARS1-KD stable cell lines. The pLVX-IRES-ZsGreen1 lentiviral vector was utilized for generating the VARS1 overexpressing plasmid. The lentiviral vectors were produced via homologous recombination cloning (Vazyme, C113) and subsequently packaged into lentiviral particles through co-transfection of three plasmids (lentiviral plasmid, psPax2, and pMD2.G) in HEK293T cells using polyethylenimine (PEI, Polysciences, 23966) at a concentration of 1 mg/mL, with a DNA:PEI ratio of 1:3 (w/w). The transfection mixture of plasmids was prepared at a molar ratio of 10:7.5:3.5 in Opti-MEM solution. The viral supernatant was harvested at 48 and 72 h post-transfection. Stable VARS1 KD or overexpressing PCa cell lines were generated through lentiviral transduction, followed by selection with puromycin (1 μg/mL for 2–4 days). The target sequences of siRNAs and shRNAs are listed in Table S12.

### Puromycin incorporation assay

To monitor the global protein synthesis, a classical puromycin incorporation assay was used [49]. Briefly, cells at ∼70% confluence were subjected to protein isolation after incubation with medium containing 1 μM puromycin dihydrochloride (Yeasen Biotech Co., 60210ES25) for 30-50 min at 37℃ and 5% CO_2_. Immunoblotting was performed with anti-puromycin antibody (Kerafast, EQ0001, dilution 1:1000). Note that the same procedure [2] was also applied to PCa cells treated with gene-specific knockdown siRNAs or chemical inhibitors.

### tRNA aminoacylation assay

Total RNA was isolated and RNA samples were resuspended in 0.3 M sodium acetate buffer (pH 4.5) containing 10 mM EDTA, followed by ethanol precipitation at −20°C. After overnight incubation, the precipitated RNA was reconstituted in 10 mM sodium acetate buffer (pH 4.5) with 1 mM EDTA. For oxidative treatment, 2 μg of RNA was incubated with 50 mM NaIO4 in darkness at room temperature for 20 min; while 2 μg control samples was incubated with 10 mM NaCl. The oxidation reaction was quenched with 100 mM glucose for 15 min. Yeast tRNA-Phe (R4018, Sigma-Aldrich) was added as an internal control prior to ethanol reprecipitation. The RNA pellets were dissolved in 50 mM Tris-HCl buffer (pH 9.0) and incubated at 37°C for 50 min. Reactions were terminated by acetate buffer-mediated precipitation. The processed RNA was subsequently ligated to 5’-adenylated DNA adapters (5’-/5rApp/TGGAATTCTCGGGTGCCAAGG/3ddC/-3’) using truncated KQ mutant T4 RNA ligase 2 (NEB, M0373) in RNase-free water at room temperature for 3 h. cDNA synthesis was performed with Induro reverse transcriptase using adaptor-complementary primers. Then, qPCR analysis was conducted with tRNA isodecoder-specific primer pairs: forward primers complementary to the 5’ end of tRNAs and reverse primers spanning the junction between tRNA 3’ ends and ligated adapters. Charge fraction values were calculated by subtracting Ct values obtained from oxidized samples (representing charged tRNA) from those of non-oxidized controls (total tRNA) using the comparative 2^−ΔΔCt^ method [50]. The primers used in this assay are listed in Table S12.

### Animal experiments and in vivo drug treatment

All experimental animals were purchased from Beijing Charles River Laboratory (Beijing, China), and housed in an SPF-grade animal facility at the School of Biomedical Sciences, Hunan University. All animal work was conducted in accordance with the protocol (HNU-IACUC-2021-101) approved by the IACUC at Hunan University. For xenograft and allograft assays, 6-7 weeks old male BALB/c-nu nude mice and 7-8 weeks old C57BL/6 mice were used, respectively. Generally, approximately 1-2 million human PCa cells and 0.5-1 million murine RM1 cells per injection were suspended in a mixture of 100 μL PBS and Matrigel (1:1), and then injected subcutaneously into male mice. Tumor volume was measured at regular intervals and the volume was calculated as (length × width^2^)/2. Finally, the mice were sacrificed and tumors were dissected. For VR treatment and drug-efficacy studies in subcutaneous tumour models, randomization was done when tumor volume reached around 100mm^3^. Animal body weight and tumor growth were measured every 3 days or twice weekly during the experiments. At the end of experiments, tumors were collected and tumor incidence, weight, and gross images were recorded. No blinding was done in the in vivo drug studies or in data analysis. For VR, three dietary regimens were used: a control (standard) diet (containing 8 g/kg valine), a 0.08% valine-restricted group (valine-free chow supplemented with 0.8 g/L valine in drinking water) and a 0.045% valine-restricted group (valine-free chow supplemented with 0.45 g/L valine in drinking water). The valine-free chow was commercially obtained from Dyets Company (Wuxi, China). For lung metastasis assay, 8 weeks old male C57BL/6 mice were intravenously injected with RM1 cells (0.3M cells per mouse) via the tail vein. After 7 days, mice were subsequently fed with an 0.8%, 0.08%, and 0.045% valine diet, respectively. After 50-60 days, mice were euthanized and the lung metastatic nodules were quantified using India ink perfusion staining. To investigate the effects of VR on late stage tumour progression in Hi-Myc PCa model, around 75 weeks old male mice (time varies due to appearance of pulpable tumors in individual mouse) were randomly assigned to two groups: a control diet and a 0.08% valine-restricted group. Body weights were measured at approximately 6-day intervals during the 50 days of treatment. The VARS1i probes were dissolved in dimethyl sulfoxide (DMSO). For in-vivo administration, VARS1i was dissolved in vehicle (10% DMSO, 20% PEG300 and 5% Tween-80 in sterile PBS) and administered via i.p. injection at indicated doses once the other day. After intended treatments, the genitourinary tract and prostate were isolated and weighed. Whole-mount prostate was subjected to H&E and IHC staining and Aperio Scanscope analysis. The animal or tumor number used for these assays were specified in the figure legends.

### Regular RNA-seq, GSEA, and GO analysis

The RNA-seq analysis was performed in collaboration with Novogene (Shanghai, China). Briefly, total RNA was extracted using TRIzol reagent and quantified using a NanoDrop spectrophotometer. RNA integrity was accessed by an Agilent 2100 Bioanalyzer. Then, the mRNAs with poly(A) tails were purified using Oligo(dT)-coated magnetic beads, followed by the first-strand cDNA synthesis with random hexamer primers. The cDNA libraries were sequenced on illumina Novaseq 6000 platform (Illumina, USA). For data analysis, we first compared clean reads with the Human reference genome (GRCh38) using Hisat2 v2.0.5, and then the DEGs were identified using DESeq2 v1.20.0 package with statistical thresholds of FDR < 0.05 and FC ≥ 1.5 or 2 (dependent on comparisons) plus a baseMean (readcounts) >10 (to remove lowly expressed genes). For pathway enrichment assay, we took a bioinformatic strategy reported by us recently [24]. GO analysis was performed on the up- or down-regulated genes using Metascape [51]. Terms with *p* < 0.01, minimum count 3, and enrichment factor >1.5 (enrichment factor is the ratio between observed count and the count expected by chance) were considered significant. Significantly enriched terms with similar descriptions and functions were further grouped into distinct biological categories (to better reflect the biology of a context) [24]. GSEA was carried out by using the curated gene sets (C2) of the Molecular Signature Database (MSigDB) v.4.059. The list of the entire detectable genes with Log2 ratios derived from each comparison was used for prerank GSEA, and we followed the standard procedure described by GSEA user guide. The FDR for GSEA is the estimated probability that a gene set with a given NES (normalized enrichment score) represents a false positive finding and an FDR < 0.25 is considered to be statistically significant.

### Polysome profiling and polysome-seq

Polysome profiling was modified from a recent report [52]. Briefly, after wash with ice-cold PBS containing 100 μg/mL cycloheximide (CHX; Amole, M4879), cells were harvested and lysed in 200 μL lysis buffer comtaining 20 mM Tris-HCl (pH 7.4), 150 mM NaCl, 5 mM MgCl2, 1% Triton X-100, 1 mM DTT, 100 μg/mL CHX, and 20 U/mL RNase inhibitor (Thermo Fisher, AM2694). RNA concentration was measured using a Nanodrop spectrometer at 260nm. Sucrose gradient solutions of 15% and 50% (wt/vol) were prepared using a polysome buffer (20 mM Tris-HCl (pH 7.4), 150 mM NaCl, 5 mM MgCl2, 1 mM DTT). The linear 15-50% sucrose gradients were generated in SW41 ultracentrifuge tubes using a BioComp Gradient Master (Biocomp Instruments, New Brunswick, Canada), followed by stabilization of the gradients at 4°C for 45 min. The lysates were then carefully loaded onto the sucrose gradients and centrifuged at 38,000 rpm for 3 h at 4°C. After centrifugation, the ribosome fractions were collected and analyzed using the Piston Gradient Fractionator^TM^ system. The monosome fraction and different polysome fractionations were combined with anhydrous ethanol at a 1:3 (v/v) ratio and incubated at −80°C overnight. The mixture was centrifuged at 12,000g and 4°C for 15 minutes, after which the supernatant was discarded. RNA-easy lysis buffer was added to the pellets and RNA of each fraction was extracted. For subsequent polysome RT-qPCR analysis [53], RNAs from extracted polysome fractions were reverse-transcribed and subjected to qPCR analysis (primers listed in Table S12). For polysome-seq, all polysome fractions were combined for RNA extraction, followed by regular RNA-seq analysis and DEGs calling as described above (conducted by Novogene company). For the purpose of analyzing transcripts at codon level, gene expression quantification was performed with RSEM (v1.3.3) [54], restricting the analysis to the longest transcript of each gene (in order to avoid redundancy in codon usage and transcript quantification analyses). Differential expression was assessed using DESeq2, applying the Benjamini–Hochberg procedure for multiple testing correction. Tanscripts with an adjusted *p* value < 0.05 and an absolute log2(fold change) > 1.5 were considered significantly differentially expressed.

### Ribo-seq and data analysis

The Ribo-seq protocol was adapted from previously reports [52, 55]. The cell lysis procedure was the same as that used in polysome profiling. A 100 μL aliquot of the cell lysate was reserved for regular RNA-seq analysis. For Ribo-seq, cell lysate was first digested with RNAse I (LGC Biosearch Technologies, E0067-10D1) at 25°C for an hour, followed by addition of 1 μL of SUPERase*In RNAse Inhibitor (Life Technologies, AM2694) to stop the reaction. Subsequently, the monosomes were collected through sucrose gradient fractionation. RNA was isolated using TRIzol (Life Technologies, 15596026) and then separated on a 16% urea-denaturing gel. After excision of ribosome-protected fragments (RPFs) around 26–34 nucleotides in length from the gel, their 3’ ends were repaired with T4 PNK (NEB, M0201) enzyme. Adapters were ligated to the RPFs using using T4 RNA Ligase 2, truncated KQ (NEB, M0242) at 22°C for 3 h. The ligated RPFs were then purified, reverse-transcribed using HiScript II Reverse Transcriptase (Vazyme, R201-01), and further purified. The cDNA was then circularized using CircLigase ssDNA ligase (LGC Biosearch Technologies, CL9025K) at 60°C for 2 hours. The product was then amplified using Q5 High-Fidelity 2X Master Mix (NEB, #M0492) and purified using AMPure XP Beads (NEB, #M0492). The final libraries were sequenced on an Illumina Novaseq 6000 platform.

For Ribo-seq data analysis, we used similar procedure as reported [52]. Cutadapt (https://cutadapt.readthedocs.io/en/stable/) was utilized to remove adaptors and random sequence, followed by quality control to trim low-quality reads. Subsequently, the contaminated rRNA and tRNA sequences were eliminated using Bowtie v1.3.1, and the remaining high-quality reads were compared to the reference human genome (Telomere-to-Telomere human genome assembly T2T-CHM13 v1.1) using STAR v2.7.3, with uniquely mapped reads of 27-35 nucleotides in length being remained for further analysis. Downstream analysis was performed with the RiboParser toolkit [56], including ribosome-protected fragment (RPF) length distribution, triplet periodicity, meta-gene analysis, and RPF quantification. To avoid redundancy in codon usage and transcript quantification analyses, only the longest transcript of each gene was retained. Codon-level translation dynamics were further evaluated using the codon pausing score, focusing on the four synonymous codons of valine (GUU, GUC, GUA, and GUG). Differentially translated genes were identified by integrating RPF quantification with RNA-seq data, restricting the analysis to genes with non-zero expression in RNA-seq, and applying the same statistical thresholds as for transcript-level analysis.

### DIA quantitative proteome

The proteomics was performed in collaboration with Novogene (Shanghai, China). The cell pellets were flash-frozen in liquid nitrogen and the company conducted protein extraction, quantification, quality control, tryptic digestion, desalting, peptide fractionation, and mass spectrometry analysis using an Orbitrap Astral high-resolution mass spectrometer. Raw data were processed using DIA-NN software for database [homo_sapiens_uniprot_2023_10_18_Swissprot.fasta (20427 sequences)] searching and peptide/protein identification. DEPs were identified based on a threshold of FC >1.2 (up-regulation) or <0.83 (down-regulation) and a statistical significance of *P* < 0.05. Human gene annotations (GTF) and coding sequences (FASTA) were obtained from Ensembl v75. The sequences underwent dual filtrations: (1) exclusion of entries lacking 5’-ATG initiation codons, and (2) removal of sequences violating triplet periodicity (length not divisible by 3) to ensure translational integrity.

Protein-gene mapping was implemented through systematic cross-referencing of Ensembl annotations. For the analysis of codon usage, the longest CDS isoform per gene was selected as the representative to quantify codon frequencies, defined as the proportion of each codon relative to the total codon count. Valine codon distributions (GTG/GTC/GTT/GTA) between DEPs (separated into up- and down-regulated groups) and non-DEGs were statistically compared using Wilcox-test or chi-square testing. Furthermore, codon usage divergence between the shVARS1 and shNC groups was determined at a significance threshold of *P* < 0.05. GO analysis of DEPs was done by Metascape as described above.

### Codon usage calculation

Codon usage was analyzed using the longest transcript of each gene downloaded from Human reference genome CHM13. Codon frequency was defined as the number of occurrences of a specific codon per 1000 coding codons in the gene set (Table S13), calculated as:

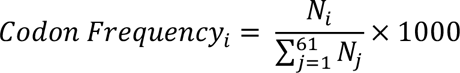

where *N_i_* represents the count of codon *i*, and the denominator is the total count of the 61 sense codons. This normalization allows comparisons across genes and gene sets.

Relative synonymous codon usage (RSCU) was calculated to assess synonymous codon preference [57]:

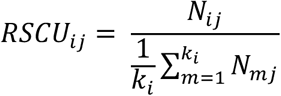

where *N_ij_* is the observed count of the *i-th* codon encoding amino acid *j*, k*_j_* is the number of synonymous codons for amino acid *j*, m = 1, 2, …, k*_j_* indexes all synonymous codons for amino acid *j*. An RSCU value of 1 indicates equal usage of synonymous codons, while values >1 suggest codon preference (Table S14).

Codon frequency and RSCU were computed using custom R scripts. The results were visualized using heatmaps and Cumulative Distribution Function (CDF) to compare codon usage patterns across gene sets. For Fig. 4B-4E and Fig. S4B, S4D, S4F-S4K, codon frequency was used to display the valine codon usage difference in genes or proteins impacted by VARS1, as this method reflects the absolute count of codon appearance and does not consider the differences in usage between synonymous codons for the same amino acid. For Fig. 4F-4G and Fig. 5A and 5H, RSCU was used to assign top genes with ‘optimal’ GTC/GTT codons, as this method reflects the relative preference for certain codons regardless of the total number of codons in a gene.

### Molecular docking

Similar to our recent report [40], a molecular docking-based virtual screening was performed against SPECS compounds (http://www.specs.net; approximately 10^6^ compounds) to identify potential ligands for human VARS1 (UniProt P26640). AlphaFold2-derived structures (VARS1: AF-P26640-F1-v4; VARS2: Q5ST30-F1-v4) were prepared in AutoDock Tools 1.5.6 with polar hydrogens and charges, while OpenBabel script-batched ligand conversion ensured PDBQT compatibility. High-throughput docking via AutoDock Vina 1.1.2 prioritized compounds by binding energy (ΔG). Protein-binding pockets were predicted using Proteinplus (https://proteins.plus/). The AutoDockTools 1.5.6 package was then utilized to generate a parameter file for the docking box, based on the predicted pocket location. Molecular docking was then performed using AutoDock Vina software, with the ligands being prepared by merging non-polar hydrogen atoms and defining rotatable bonds. Hydrogen bond interactions were then captured using PyMOL, and a two-dimensional interaction diagram was obtained from the ProteinPlus website. The coordinates of the active pocket of the VARS1 protein molecule are set to: centre_x = 10.137, centre_y = 1.818, centre_z = −11.350. The size parameters are set to 114 for size_x, 72 for size_y, and 68 for size_z. In order to increase the docking accuracy, the value of exhaustiveness was set to 16. For Vina docking, the default parameters were used if it was not specified. The best-scoring pose as judged by the Vina docking score was chosen and visually analyzed using PyMoL 1.7.6 software (www.pymol.org).

### In vitro fluorescence titration experiment

The CDS of human VARS1 was cloned into pET-28a plasmid and used for subsequent prokaryotic expression. 100 µL of VARS1 proteins (10 µM) was added to the 96-well plate, followed by the addition of an equal volume of Buffer containing compounds at the final dose of 0.01, 0.02, 2, 4, 6, 8 and 10 μM respectively. The plate was gently blown and incubated at 4 °C for 10 minutes, then the fluorescence value of each well was measured under excitation light at 280 nm. The fluorescence value of the buffer wells with compounds alone were used to exclude the optical interference of the compound. The resultant data were then examined for evidence of a decrease in fluorescence intensity with an increase in compound amount, thus allowing the dose-effect relationship to be determined. The equilibrium dissociation constant (Kd) between a protein and a small molecule was calculated using Origin 2024, representing the concentration at which half the protein is bound. This reflects the compound’s affinity for the target, and a smaller value indicates a stronger affinity.

### Statistical analysis

Statistical analyses were performed using GraphPad Prism 9 software or using R. In general, unpaired two-tailed Student’s t-test or Wilcoxon’s signed-rank tests, and χ^2^-test were used to calculate the statistical significance between pair comparisons depending on the data type. Multigroup comparisons were analyzed using one-way ANOVA. Correlation analyses were conducted using Pearson or Spearman tests. *P* < 0.05 is considered statistically significant. The specific statistical methods used for each experiment are detailed in the text or figure legends.

## Acknowledgements

This work was supported by grants from the National Natural Science Foundation of China 32470766 (to D.Z.), 32470073 (to Z.Z.) and 32500462 (to C.Z.), National Key Research and Development Program of China 2024YFC2510500 and Natural Science Foundation of Shandong Province ZR2024MH065 (to B.Z.), the National Clinical key Discipline Construction Funds CZXM-ZK-47 and Jiangsu Provincial Key Discipline and Laboratory Construction Funds of Urology 2023YXZDXK02 (to B.X.), Guangdong Basic and Applied Basic Research Foundation 2025A1515012400 (to D.Z.), and the Fundamental Research Funds for the Central Universities 531119200130 (to D.Z.). C.Z. was also supported, in part, by the China Postdoctoral Science Foundation (2024M750867). We thank Dr. Zhenfei Li and Dong Gao (Shanghai Institute of Biochemistry and Cell Biology) for sharing reagents. We apologize to the colleagues whose work was not cited due to space constraint.

## Author contributions

D.Z. and Z.Z. conceived and designed the study. Q.H., B.Z., Z.Z. and D.Z. interpreted the data. D.Z. wrote and finalized the manuscript with help from Q.H. and S.R. Q.H., S.R., C.Z. and D.Z. conducted the bioinformatic analysis. Q.H., R.H. and Y.L. performed most experiments under the supervision of D.Z. M.X. and B.Z. carried out the virtual drug screen and molecular docking. Y.Z and L.H. assisted with in vitro cell-based assays and in vivo animal studies. B.X. and W.L. provided clinical samples and IHC analysis. All authors read and approved the manuscript.

## Data Availability Statement

The data that support the findings of this study have been deposited in the Gene Expression Omnibus (GEO) under the accession codes GSE307636 (Ribo-seq), GSE307637 (RNA-seq of VARS1 KD), GSE307495 (Polysome-seq), and GSE306670 (RNA-seq of valine restriction). Mass spectrometry proteomics data have been deposited to the ProteomeXchange Consortium via the iProX partner repository with the accession code PXD068236. All other data supporting the results are available within the article and its Supplementary Information files and from the corresponding authors upon request.

## Supplementary Tables

Table S1. List of 105 genes involved in tRNA biogenesis and function.

Table S2. List of differentially expressed TBGs in indicated comparisons.

Table S3. GSEA of genes coexpressed with VARS1 in both pri-PCa and CRPC cohorts.

Table S4. List of DEGs and GSEA of genes downregulated in PC3 cells upon VARS1 KD.

Table S5. List of DEGs identified in polysome-seq in PC3 cells with or without VARS1 KD.

Table S6. List of DEGs identified in Ribo-seq in PC3 cells upon VARS1 depletion.

Table S7. List of DEPs identified in proteomes of PC3 cells with or without VARS1 KD.

Table S8. Gene list of VARS1-translational signature.

Table S9. List of DEPs in PC3 cells cultured in low vs. high valine medium.

Table S10. List of DEGs in PC3 cells cultured in low vs. high valine medium.

Table S11. List of Specs compounds identified in virtual drug screen.

Table S12. List of primers, siRNAs, shRNAs and antibodies used in this study

Table S13. Valine codon frequency calculated for each human gene

Table S14. Valine RSCU score calculated for each human gene

## REFERENCES

[1] R.L. Siegel, A.N. Giaquinto, A. Jemal, Cancer statistics, 2024, CA: a cancer journal for clinicians, 74 (2024) 12–49.

[2] C. Zou, W. Li, Y. Zhang, N. Feng, S. Chen, L. Yan, Q. He, K. Wang, W. Li, Y. Li, Y. Wang, B. Xu, D. Zhang, Identification of an anaplastic subtype of prostate cancer amenable to therapies targeting SP1 or translation elongation, Sci Adv, 10 (2024) eadm7098.

[3] P.A. Watson, V.K. Arora, C.L. Sawyers, Emerging mechanisms of resistance to androgen receptor inhibitors in prostate cancer, Nature reviews. Cancer, 15 (2015) 701–711.

[4] Q. Li, Q. Deng, H.P. Chao, X. Liu, Y. Lu, K. Lin, B. Liu, G.W. Tang, D. Zhang, A. Tracz, C. Jeter, K. Rycaj, T. Calhoun-Davis, J. Huang, M.A. Rubin, H. Beltran, J. Shen, G. Chatta, I. Puzanov, J.L. Mohler, J. Wang, R. Zhao, J. Kirk, X. Chen, D.G. Tang, Linking prostate cancer cell AR heterogeneity to distinct castration and enzalutamide responses, Nature communications, 9 (2018) 3600.

[5] C. Zou, Q. He, Y. Feng, M. Chen, D. Zhang, A m(6)Avalue predictive of prostate cancer stemness, tumor immune landscape and immunotherapy response, NAR cancer, 4 (2022) zcac010.

[6] M. Cai, X.L. Song, X.A. Li, M. Chen, J. Guo, D.H. Yang, Z. Chen, S.C. Zhao, Current therapy and drug resistance in metastatic castration-resistant prostate cancer, Drug Resist Updat, 68 (2023) 100962.

[7] N. Cancer Genome Atlas Research, The Molecular Taxonomy of Primary Prostate Cancer, Cell, 163(2015) 1011-1025.

[8] D. Robinson, E.M. Van Allen, Y.M. Wu, N. Schultz, R.J. Lonigro, J.M. Mosquera, B. Montgomery, M.E. Taplin, C.C. Pritchard, G. Attard, H. Beltran, W. Abida, R.K. Bradley, J. Vinson, X. Cao, P. Vats, L.P. Kunju, M. Hussain, F.Y. Feng, S.A. Tomlins, K.A. Cooney, D.C. Smith, C. Brennan, J. Siddiqui, R. Mehra, Y. Chen, D.E. Rathkopf, M.J. Morris, S.B. Solomon, J.C. Durack, V.E. Reuter, A. Gopalan, J. Gao, M. Loda, R.T. Lis, M. Bowden, S.P. Balk, G. Gaviola, C. Sougnez, M. Gupta, E.Y. Yu, E.A. Mostaghel, H.H. Cheng, H. Mulcahy, L.D. True, S.R. Plymate, H. Dvinge, R. Ferraldeschi, P. Flohr, S. Miranda, Z. Zafeiriou, N. Tunariu, J. Mateo, R. Perez-Lopez, F. Demichelis, B.D. Robinson, M. Schiffman, D.M. Nanus, S.T. Tagawa, A. Sigaras, K.W. Eng, O. Elemento, A. Sboner, E.I. Heath, H.I. Scher, K.J. Pienta, P. Kantoff, J.S. de Bono, M.A. Rubin, P.S. Nelson, L.A. Garraway, C.L. Sawyers, A.M. Chinnaiyan, Integrative clinical genomics of advanced prostate cancer, Cell, 161 (2015) 1215–1228.

[9] A. Sinha, V. Huang, J. Livingstone, J. Wang, N.S. Fox, N. Kurganovs, V. Ignatchenko, K. Fritsch, N. Donmez, L.E. Heisler, Y.J. Shiah, C.Q. Yao, J.A. Alfaro, S. Volik, A. Lapuk, M. Fraser, K. Kron, A. Murison, M. Lupien, C. Sahinalp, C.C. Collins, B. Tetu, M. Masoomian, D.M. Berman, T. van der Kwast, R.G. Bristow, T. Kislinger, P.C. Boutros, The Proteogenomic Landscape of Curable Prostate Cancer, Cancer cell, 35 (2019) 414–427 e416.

[10] B. Dong, J.Y. Xu, Y. Huang, J. Guo, Q. Dong, Y. Wang, N. Li, Q. Liu, M. Zhang, Q. Pan, H. Wang, J. Jiang, B. Chen, D. Shen, Y. Ma, L. Zhai, J. Zhang, J. Li, W. Xue, M. Tan, J. Qin, Integrative proteogenomic profiling of high-risk prostate cancer samples from Chinese patients indicates metabolic vulnerabilities and diagnostic biomarkers, Nat Cancer, 5 (2024) 1427–1447.

[11] R. Sun, J. A, H. Yu, Y. Wang, M. He, L. Tan, H. Cheng, J. Zhang, Y. Wang, X. Sun, M. Lyu, M. Qu, L. Huang, Z. Li, W. Zhang, K. Ma, Z. Dong, W. Ge, Y. Zhang, X. Ding, B. Yang, J. Hou, C. Xu, L. Wang, Y. Zhu, T. Guo, X. Gao, C. Yang, Proteomic landscape profiling of primary prostate cancer reveals a 16-protein panel for prognosis prediction, Cell Rep Med, 5 (2024) 101679.

[12] Z. Wang, H. Yu, W. Bao, M. Qu, Y. Wang, L. Zhang, X. Liu, C. Liu, M. He, J. Li, Z. Dong, Y. Zhang, B. Yang, J. Hou, C. Xu, L. Wang, X. Li, X. Gao, C. Yang, Proteomic and phosphoproteomic landscape of localized prostate cancer unveils distinct molecular subtypes and insights into precision therapeutics, Proceedings of the National Academy of Sciences of the United States of America, 121 (2024) e2402741121.

[13] L. Fabbri, A. Chakraborty, C. Robert, S. Vagner, The plasticity of mRNA translation during cancer progression and therapy resistance, Nature reviews. Cancer, 21 (2021) 558–577.

[14] P. Thandapani, A. Kloetgen, M.T. Witkowski, C. Glytsou, A.K. Lee, E. Wang, J. Wang, S.E. LeBoeuf, K. Avrampou, T. Papagiannakopoulos, A. Tsirigos, I. Aifantis, Valine tRNA levels and availability regulate complex I assembly in leukaemia, Nature, 601 (2022) 428–433.

[15] Y. Liu, J.L. Horn, K. Banda, A.Z. Goodman, Y. Lim, S. Jana, S. Arora, A.A. Germanos, L. Wen, W.R. Hardin, Y.C. Yang, I.M. Coleman, R.G. Tharakan, E.Y. Cai, T. Uo, S.P.S. Pillai, E. Corey, C. Morrissey, Y. Chen, B.S. Carver, S.R. Plymate, S. Beronja, P.S. Nelson, A.C. Hsieh, The androgen receptor regulates a druggable translational regulon in advanced prostate cancer, Sci Transl Med, 11 (2019).

[16] A.M. Pinzaru, S.F. Tavazoie, Transfer RNAs as dynamic and critical regulators of cancer progression, Nature reviews. Cancer, 23 (2023) 746–761.

[17] I. Yoon, U. Kim, J. Choi, S. Kim, Disease association and therapeutic routes of aminoacyl-tRNA synthetases, Trends in molecular medicine, 30 (2024) 89–105.

[18] M.C. Passarelli, A.M. Pinzaru, H. Asgharian, M.V. Liberti, S. Heissel, H. Molina, H. Goodarzi, S.F. Tavazoie, Leucyl-tRNA synthetase is a tumour suppressor in breast cancer and regulates codon-dependent translation dynamics, Nature cell biology, 24 (2022) 307–315.

[19] K. Kurata, A. James-Bott, M.A. Tye, L. Yamamoto, M.K. Samur, Y.T. Tai, J. Dunford, C. Johansson, F. Senbabaoglu, M. Philpott, C. Palmer, K. Ramasamy, S. Gooding, M. Smilova, G. Gaeta, M. Guo, J.C. Christianson, N.C. Payne, K. Singh, K. Karagoz, M.E. Stokes, M. Ortiz, P. Hagner, A. Thakurta, A. Cribbs, R. Mazitschek, T. Hideshima, K.C. Anderson, U. Oppermann, Prolyl-tRNA synthetase as a novel therapeutic target in multiple myeloma, Blood Cancer J, 13 (2023) 12.

[20] D.G. Song, D. Kim, J.W. Jung, S.H. Nam, J.E. Kim, H.J. Kim, J.H. Kim, S.J. Lee, C.H. Pan, S. Kim, J.W. Lee, Glutamyl-prolyl-tRNA synthetase induces fibrotic extracellular matrix via both transcriptional and translational mechanisms, FASEB J, 33 (2019) 4341–4354.

[21] J. Gao, B.A. Aksoy, U. Dogrusoz, G. Dresdner, B. Gross, S.O. Sumer, Y. Sun, A. Jacobsen, R. Sinha, E. Larsson, E. Cerami, C. Sander, N. Schultz, Integrative analysis of complex cancer genomics and clinical profiles using the cBioPortal, Sci Signal, 6 (2013) pl1.

[22] J.Z. Ruidong Li, Wei-De Zhong, Zhenyu Jia, PCaDB: A comprehensive and interactive database for transcriptomes from prostate cancer population cohorts, bioRxiv, (2022).

[23] W. Abida, J. Cyrta, G. Heller, D. Prandi, J. Armenia, I. Coleman, M. Cieslik, M. Benelli, D. Robinson, E.M. Van Allen, A. Sboner, T. Fedrizzi, J.M. Mosquera, B.D. Robinson, N. De Sarkar, L.P. Kunju, S. Tomlins, Y.M. Wu, D. Nava Rodrigues, M. Loda, A. Gopalan, V.E. Reuter, C.C. Pritchard, J. Mateo, D. Bianchini, S. Miranda, S. Carreira, P. Rescigno, J. Filipenko, J. Vinson, R.B. Montgomery, H. Beltran, E.I. Heath, H.I. Scher, P.W. Kantoff, M.E. Taplin, N. Schultz, J.S. deBono, F. Demichelis, P.S. Nelson, M.A. Rubin, A.M. Chinnaiyan, C.L. Sawyers, Genomic correlates of clinical outcome in advanced prostate cancer, Proceedings of the National Academy of Sciences of the United States of America, 116 (2019) 11428–11436.

[24] D. Zhang, Q. Hu, X. Liu, Y. Ji, H.P. Chao, Y. Liu, A. Tracz, J. Kirk, S. Buonamici, P. Zhu, J. Wang, S. Liu, D.G. Tang, Intron retention is a hallmark and spliceosome represents a therapeutic vulnerability in aggressive prostate cancer, Nature communications, 11 (2020) 2089.

[25] V. Reddy, A. Iskander, C. Hwang, G. Divine, M. Menon, E.R. Barrack, G.P. Reddy, S.H. Kim, Castration-resistant prostate cancer: Androgen receptor inactivation induces telomere DNA damage, and damage response inhibition leads to cell death, PloS one, 14 (2019) e0211090.

[26] S. Tai, Y. Sun, J.M. Squires, H. Zhang, W.K. Oh, C.Z. Liang, J. Huang, PC3 is a cell line characteristic of prostatic small cell carcinoma, The Prostate, 71 (2011) 1668–1679.

[27] N.H. Kwon, P.L. Fox, S. Kim, Aminoacyl-tRNA synthetases as therapeutic targets, Nature reviews. Drug discovery, 18 (2019) 629–650.

[28] M. Guo, X.L. Yang, P. Schimmel, New functions of aminoacyl-tRNA synthetases beyond translation, Nature reviews. Molecular cell biology, 11 (2010) 668–674.

[29] X.D. He, W. Gong, J.N. Zhang, J. Nie, C.F. Yao, F.S. Guo, Y. Lin, X.H. Wu, F. Li, J. Li, W.C. Sun, E.D. Wang, Y.P. An, H.R. Tang, G.Q. Yan, P.Y. Yang, Y. Wei, Y.Z. Mao, P.C. Lin, J.Y. Zhao, Y. Xu, W. Xu, S.M. Zhao, Sensing and Transmitting Intracellular Amino Acid Signals through Reversible Lysine Aminoacylations, Cell metabolism, 27 (2018) 151–166 e156.

[30] J. Ju, H. Zhang, M. Lin, Z. Yan, L. An, Z. Cao, D. Geng, J. Yue, Y. Tang, L. Tian, F. Chen, Y. Han, W. Wang, S. Zhao, S. Jiao, Z. Zhou, The alanyl-tRNA synthetase AARS1 moonlights as a lactyltransferase to promote YAP signaling in gastric cancer, The Journal of clinical investigation, 134 (2024).

[31] H. Li, C. Liu, R. Li, L. Zhou, Y. Ran, Q. Yang, H. Huang, H. Lu, H. Song, B. Yang, H. Ru, S. Lin, L. Zhang, AARS1 and AARS2 sense L-lactate to regulate cGAS as global lysine lactyltransferases, Nature, 634 (2024) 1229–1237.

[32] Y. Sung, I. Yoon, J.M. Han, S. Kim, Functional and pathologic association of aminoacyl-tRNA synthetases with cancer, Experimental & molecular medicine, 54 (2022) 553–566.

[33] N. El-Hachem, M. Leclercq, M. Susaeta Ruiz, R. Vanleyssem, K. Shostak, P.R. Körner, C. Capron, L. Martin-Morales, P. Roncarati, A. Lavergne, A. Blomme, S. Turchetto, E. Goffin, P. Thandapani, I. Tarassov, L. Nguyen, B. Pirotte, A. Chariot, J.C. Marine, M. Herfs, F. Rapino, R. Agami, P. Close, Valine aminoacyl-tRNA synthetase promotes therapy resistance in melanoma, Nature cell biology, 26 (2024) 1154–1164.

[34] D. Meyer, J. Kames, H. Bar, A.A. Komar, A. Alexaki, J. Ibla, R.C. Hunt, L.V. Santana-Quintero, A. Golikov, M. DiCuccio, C. Kimchi-Sarfaty, Distinct signatures of codon and codon pair usage in 32 primary tumor types in the novel database CancerCoCoPUTs for cancer-specific codon usage, Genome Med, 13 (2021) 122.

[35] M. Chen, C. Zou, Y. Tian, W. Li, Y. Li, D. Zhang, An integrated ceRNA network identifies miR-375 as an upregulated miRNA playing a tumor suppressive role in aggressive prostate cancer, Oncogene, 43 (2024) 1594–1607.

[36] B. van de Kooij, F.J. van der Wal, M.B. Rother, W.W. Wiegant, P. Creixell, M. Stout, B.A. Joughin, J. Vornberger, M. Altmeyer, M. van Vugt, M.B. Yaffe, H. van Attikum, The Fanconi anemia core complex promotes CtIP-dependent end resection to drive homologous recombination at DNA double-strand breaks, Nature communications, 15 (2024) 7076.

[37] M. Pilkinton, R. Sandoval, K. Barrett, X. Tian, O.R. Colamonici, Mip/LIN-9 can inhibit cell proliferation independent of the pocket proteins, Blood Cells Mol Dis, 39 (2007) 272–277.

[38] K.M. Janisch, V.M. Vock, M.S. Fleming, A. Shrestha, C.M. Grimsley-Myers, B.A. Rasoul, S.A. Neale, T.D. Cupp, J.M. Kinchen, K.F. Liem, Jr., N.D. Dwyer, The vertebrate-specific Kinesin-6, Kif20b, is required for normal cytokinesis of polarized cortical stem cells and cerebral cortex size, Development, 140 (2013) 4672–4682.

[39] C. Zhang, M. Kuang, M. Li, L. Feng, K. Zhang, S. Cheng, SMC4, which is essentially involved in lung development, is associated with lung adenocarcinoma progression, Sci Rep, 6 (2016) 34508.

[40] P. Feng, D. Chen, X. Wang, Y. Li, Z. Li, B. Li, Y. Zhang, W. Li, J. Zhang, J. Ye, B. Zhao, J. Li, C. Ji, Inhibition of the m(6)A reader IGF2BP2 as a strategy against T-cell acute lymphoblastic leukemia, Leukemia, 36 (2022) 2180–2188.

[41] K. Ellwood-Yen, T.G. Graeber, J. Wongvipat, M.L. Iruela-Arispe, J. Zhang, R. Matusik, G.V. Thomas, C.L. Sawyers, Myc-driven murine prostate cancer shares molecular features with human prostate tumors, Cancer cell, 4 (2003) 223–238.

[42] D. Zhang, C. Jeter, S. Gong, A. Tracz, Y. Lu, J. Shen, D.G. Tang, Histone 2B-GFP Label-Retaining Prostate Luminal Cells Possess Progenitor Cell Properties and Are Intrinsically Resistant to Castration, Stem Cell Reports, 10 (2018) 228–242.

[43] J. Wang, I. Vallee, A. Dutta, Y. Wang, Z. Mo, Z. Liu, H. Cui, A.I. Su, X.L. Yang, Multi-Omics Database Analysis of Aminoacyl-tRNA Synthetases in Cancer, Genes (Basel), 11 (2020).

[44] S.H. Kim, S. Bae, M. Song, Recent Development of Aminoacyl-tRNA Synthetase Inhibitors for Human Diseases: A Future Perspective, Biomolecules, 10 (2020).

[45] A.H. Diacon, C.E. Barry, 3rd, A. Carlton, R.Y. Chen, M. Davies, V. de Jager, K. Fletcher, G. Koh, I. Kontsevaya, J. Heyckendorf, C. Lange, M. Reimann, S.L. Penman, R. Scott, G. Maher-Edwards, S. Tiberi, G. Vlasakakis, C.M. Upton, D.B. Aguirre, A first-in-class leucyl-tRNA synthetase inhibitor, ganfeborole, for rifampicin-susceptible tuberculosis: a phase 2a open-label, randomized trial, Nat Med, 30 (2024) 896–904.

[46] T.L. Keller, D. Zocco, M.S. Sundrud, M. Hendrick, M. Edenius, J. Yum, Y.J. Kim, H.K. Lee, J.F. Cortese, D.F. Wirth, J.D. Dignam, A. Rao, C.Y. Yeo, R. Mazitschek, M. Whitman, Halofuginone and other febrifugine derivatives inhibit prolyl-tRNA synthetase, Nature chemical biology, 8 (2012) 311–317.

[47] H. Bharathkumar, C.D. Mohan, S. Rangappa, T. Kang, H.K. Keerthy, J.E. Fuchs, N.H. Kwon, A. Bender, S. Kim, Basappa, K.S. Rangappa, Screening of quinoline, 1,3-benzoxazine, and 1,3-oxazine-based small molecules against isolated methionyl-tRNA synthetase and A549 and HCT116 cancer cells including an in silico binding mode analysis, Org Biomol Chem, 13 (2015) 9381–9387.

[48] Z. Li, Y. Feng, H. Han, X. Jiang, W. Chen, X. Ma, Y. Mei, D. Yuan, D. Zhang, J. Shi, A Stapled Peptide Inhibitor Targeting the Binding Interface of N6-Adenosine-Methyltransferase Subunits METTL3 and METTL14 for Cancer Therapy, Angew Chem Int Ed Engl, 63 (2024) e202402611.

[49] J. Liu, Y. Xu, D. Stoleru, A. Salic, Imaging protein synthesis in cells and tissues with an alkyne analog of puromycin, Proceedings of the National Academy of Sciences of the United States of America, 109 (2012) 413–418.

[50] D. Zhang, D. Park, Y. Zhong, Y. Lu, K. Rycaj, S. Gong, X. Chen, X. Liu, H.P. Chao, P. Whitney, T. Calhoun-Davis, Y. Takata, J. Shen, V.R. Iyer, D.G. Tang, Stem cell and neurogenic gene-expression profiles link prostate basal cells to aggressive prostate cancer, Nature communications, 7 (2016) 10798.

[51] Y. Zhou, B. Zhou, L. Pache, M. Chang, A.H. Khodabakhshi, O. Tanaseichuk, C. Benner, S.K. Chanda, Metascape provides a biologist-oriented resource for the analysis of systems-level datasets, Nature communications, 10 (2019) 1523.

[52] R. He, Z. Lv, Y. Li, S. Ren, J. Cao, J. Zhu, X. Zhang, H. Wu, L. Wan, J. Tang, S. Xu, X.L. Chen, Z. Zhou, tRNA-m(1)A methylation controls the infection of Magnaporthe oryzae by supporting ergosterol biosynthesis, Developmental cell, 59 (2024) 2931–2946 e2937.

[53] A.C. Panda, J.L. Martindale, M. Gorospe, Polysome Fractionation to Analyze mRNA Distribution Profiles, Bio Protoc, 7 (2017).

[54] B. Li, C.N. Dewey, RSEM: accurate transcript quantification from RNA-Seq data with or without a reference genome, BMC bioinformatics, 12 (2011) 323.

[55] N.J. McGlincy, N.T. Ingolia, Transcriptome-wide measurement of translation by ribosome profiling, Methods, 126 (2017) 112–129.

[56] S. Ren, Y. Li, Z. Zhou, RiboParser/RiboShiny: an integrated platform for comprehensive analysis and visualization of Ribo-seq data, J Genet Genomics, (2025).

[57] P.M. Sharp, W.H. Li, The codon Adaptation Index--a measure of directional synonymous codon usage bias, and its potential applications, Nucleic Acids Res, 15 (1987) 1281–1295.

